# Inference of recent admixture using genotype data

**DOI:** 10.1101/2020.09.16.300640

**Authors:** Peter Pfaffelhuber, Elisabeth Sester-Huss, Franz Baumdicker, Jana Naue, Sabine Lutz-Bonengel, Fabian Staubach

## Abstract

The inference of biogeographic ancestry (BGA) has become a focus of forensic genetics. Misinference of BGA can have profound unwanted consequences for investigations and society. We show that recent admixture can lead to misclassification and erroneous inference of ancestry proportions, using state of the art analysis tools with (i) simulations, (ii) 1000 genomes project data, and (iii) two individuals analyzed using the ForenSeq DNA Signature Prep Kit. Subsequently, we extend existing tools for estimation of individual ancestry (IA) by allowing for different IA in both parents, leading to estimates of parental individual ancestry (PIA), and a statistical test for recent admixture. Estimation of PIA outperforms IA in most scenarios of recent admixture. Furthermore, additional information about parental ancestry can be acquired with PIA that may guide casework.

**Highlights:**

- We improve statistical methods as used in STRUCTURE and ADMIXTURE for Biogeographical Ancestry (BGA) inference to account for recent admixture, i.e. different admixture of both parents.
- The resulting recent admixture model has a higher accuracy in estimating individual admixture in most cases.
- We give a likelihood ratio test for recent admixture, which is both highly specific and sensitive for recent admixture.
- We find evidence of recent admixture in the 1000 genomes dataset.
- The self-report on recent admixture of two self-sequenced samples was only confirmed in one case.

## 1 Introduction

Inference of the biogeographical ancestry of a trace or an unknown body, using genetic markers, is a focus of recent forensic genetics research (see e.g. [1, 2, 3, 4]). Misclassification and erroneous inference of ancestry proportions can mislead casework and result in unwanted societal consequences [5]. Therefore, discovering potential error sources and improving methods for BGA is essential for successful and responsible application of the technology.

Throughout, we assume to have a single-source trace, i.e. genetic autosomal material from a single person. The inference of BGA of that person can mean one of two things: (i) The trace is classified into one (and only one) of several groups of different origin (e.g. Africa, Europe, East Asia, Native America, South-East Asia and Oceania; see e.g. [6, 7]). (ii) It is assumed that the single-source trace consists of a mixture of ancestral genetic material originating in several (continental) groups. For the inference of such admixture proportions of individual ancestry (IA), structure [4] and the faster admixture [3] have become the de-facto standards; see also [5] for the same model. In this manuscript, we are mainly interested in (ii).

To understand how recent admixture can cause errors in BGA inference, consider a recently admixed individual, i.e. the continental BGA of both parents differs. Since methods used in BGA classification [6, 7] can only result in single population/class label, the possibility of recent admixture is usually not even implemented by such methods, making results on classification of recently admixed individuals hard to interpret. In mixed membership models as implemented in STRUCTURE or ADMIXTURE for estimating continental (or other scales of geography) IA of a trace, the genome is thought to be a mosaic of stones of different (continental) origin. The geographic distribution of the mosaic stones (alleles) is given by the IA. Here, a main assumption is that all alleles have the same chance (given by the IA) to stem from one of the continents. However, if the BGA of the two parents differs, the chance to encounter two different alleles at a locus increases due to population differentiation, e.g. between continents [11]. This is expected to lead to a genome wide increase in heterozygosity. The violation of the assumption of equal chances for homologous allelic states could potentially lead to misinferences. Given that recent admixture is a common issue in the light of increased human mobility, currently and in the past decades, such misinference could be a common error source in forensic genetic analyses. Therefore, a more comprehensive understanding of the consequences and frequency of recent admixture in forensic analyses is needed.

For forensic applications the approaches taken by Zou et al. (2015) [12] and Pei et al. (2020) [13] to infer recent admixture are often not feasible because they rely on the inference of phase along the chromosome with dense marker sets. Furthermore, these approaches are computationally demanding. Crouch and Weale (2012) developed the LEAPFrOG algorithm that can be used with the limited marker density of most forensic applications [14]. These authors inferred the parental IA with a maximum likelihood approach and applied their method to simulated and forensically relevant datasets with a focus on European/African admixture. To identify recently admixed individuals in forensic samples, an excess of heterozygous sites was used in statistical tests [15]. Tvedebrink et al. (2018) and Tvedebrink and Eriksen (2019) developed likelihood-ratio tests for the null-hypothesis of non-admixture and recent (first generation) admixture, respectively, versus the alternative that the studied sample is not represented in the reference database [16, 17].

What is currently missing is an approach to test for recent admixture in a trace, where the null-hypothesis is that both parents have the same IA. The alternative hypothesis, called the recent-admixture model below, would be that the studied sample has parents of different IA and therefore shows recent-admixture of populations within the reference database. If such an approach also identifies the IAs of the parents of an admixed individual, this might be informative for casework. Moreover, for a better understanding of the potential misinference in forensic applications, simulations should be based on the most realistic human population genetic models and include the analysis of recent methods and marker sets. Given global mobility of humans, global sample collections should be included in the analyses.

Our goals were to identify and quantify potential errors that result from recent admixture in standard methods for BGA inference and to develop a statistical test for recent admixture vs the null hypothesis of non-admixture. To this end, we developed a method for the inference of admixture proportions of both parents (parental individual ancestry, PIA). We use this method to (i) assign IA to the parents, (ii) improve current methods of ancestry inference, (iii) perform a likelihood ratio test to identify recently admixed individuals, including the report of a *p*-value for the null hypothesis of no recent admixture. For assessing our method and also to assess the misinference of BGA with standard methods in the context of recent admixture, we leverage population genetic simulations including the most realistic human population genetic scenarios. Furthermore, we analyse a global dataset from the 1000 genomes project, and two additional samples using the ForenSeq Signature Prep Kit on a MiSeq FGx (Verogen).

## 2 Materials and Methods

### 2.1 The admixture and recent admixture model

We start by briefly recalling the admixture model, which is the basis for the widely used sofware structure [4], admixture [3], frappe [5] and the R-package LEA [18]. Afterwards, we introduce a new model, called the recent-admixture model. More details on the derivations in the admixture and recent-admixture model can be found in Section S1 in the SI. Moreover, the implementation of our methods can be downloaded from https://github.com/pfaffelh/recent-admixture.

Here comes an informal description of the models and the inference procedure. For both, the admixture and recent-admixture model, we assume to have a reference database of *M* bi-allelic markers from *K* populations. However, from this reference database, we only need to know allele frequencies, i.e. *p_mk_* is the frequency of allele 1 (or reference allele) at marker *m* in population *k* for all *m* = 1,…, *M* and *k* = 1,…, *K*. We have a trace with *G_m_* ∈ {0, 1, 2} copies of allele 1 at marker *m* for *m* = 1,…, *M* and we denote the full trace by *G* = (*G_m_*)_*m*=1,…, *M*_. The admixture model assumes that a fraction *q_k_* (to be estimated) of the genome of the trace comes from population *k* for *k* = 1,…, *K*. We will denote *q* = (*q*_1_,…, *q_K_*) the individual admixture (IA) of the trace. Estimation of *q* is done via maximum likelihood, where linkage equilibrium within populations of all markers is assumed; see (S3) for the corresponding log-likelihood-function. In the recent-admixture model, we assume that the trace originates from two parents which have (potentially) different IAs, *q^M^* and *q^P^*, denoted parental individual admixture (PIA) of the trace. The upshot is that the two allelic copies of the trace do not segregate independently, since each parent gave one copy to their child causing the trace. Again, this leads to a log-likelihood; see (S6). Since the two models are nested (the admixture model is the special case *q^M^* = *q^P^* in the recent-admixture model), we can subsequently use the likelihood-ratio-test framework for a formal test. While superscripts *M* and *P* indicate maternal and paternal ancestry, we note that the estimation of *q^M^* and *q^P^* separately is not possible since we are only dealing with autosomal markers. In terms of our method, the likelihood is symmetric in *q^M^* and *q^P^*, so we can only estimate IA of the parents irrespective which is paternal and which is maternal.

Briefly, we collect the main assumptions of both models, based exclusively on autosomal data: The ancestral populations are assumed to be homogeneous in terms of allele frequencies. Markers are assumed to be in linkage equilibrium within populations. For the trace in the admixture model (its parents in the recent-admixture model), we assume that each allele (from the parents of the trace) has the same independent chance to come from one of the ancestral groups.

We will assume throughout that the allele frequencies *p_mk_* are given and will not be changed by analysing the trace. This is important since in currently used software STRUCTURE, ADMIXUTURE and FRAPPE, mostly in non-forensic use, it is frequently the case that many new individuals are studied, and allele frequencies are updated. For forensic use, when analysing several traces at once, this would imply that the results for the ancestry of trace 1 depend not only on the reference data, but also on the data for traces 2, 3,… which seems inappropriate. Hence, we do not make the computational overload of updating allele frequencies, which would also lead to increased runtimes. Instead, we take the allele frequencies as given in the reference database. This approach has also been used in early papers such as [1] and [2].

### 2.2 Obtaining admixed individuals in silico

In the sequel, we call an individual anciently admixed if its IA satisfies *q* = *q^M^* = *q^P^*, and recently admixed if *q^M^* ≠ *q^P^*. If we are given allele frequencies *p_mk_* for all markers *m* = 1,…, *M* in all populations *k* = 1,…, *K*, we can obtain an anciently admixed individual with IA given by q by choosing the genome *G* = (*G_m_*)_*m*=1,…, *M*_ with independent markers with *G_m_* ~ *B*(2, *β_m_*(*q*)), the binomial distribution with two trials and success probability 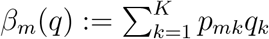. In addition, if we are given two genomes 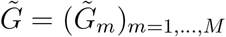 and 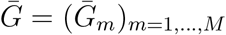 with different IA, we can obtain a recently admixed genome *G* = (*G_m_*)_*m*=1…, *M*_ by setting *G_m_* = *X_m_*+*Y_m_* with 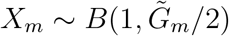 and 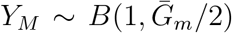. When iterating this procedure, we can also model second-generation admixed individuals etc. in silico.

Consider data in a reference database (e.g. a subset of the 1000 genomes dataset, see Section 2.5), consisting of Africans (AFR), East-Asians (EAS), Europeans (EUR) and South-East-Asians (SAS). All cases for second generation admixed individuals fall into one of seven categories. Writing up the ancestries of the four grandparents *Mother of mother/father of mother×mother of father/father of father*, we have the following distinguishable cases for second generation admixed individuals (the full list of all resulting 55 cases is given in Section S2.2 in the SI; note that [21] come up with only 35 cases, since they do not distinguish between maternal and paternal ancestry, e.g. they count AFR/AFR×EAS/EAS and AFR/EAS×AFR/EAS as one case):

A. 4 non-admixed cases, e.g. AFR/AFR× AFR/AFR;
B. 6 admixed cases with admixture ratio 50:50, where both parents are non-admixed, e.g. AFR/AFR×EAS/EAS;
C. 6 admixed cases with admixture ratio 50:50, where both parents are admixed, e.g. AFR/EAS×AFR/EAS;
D. 12 admixed cases with admixture ratio 75:25, e.g. AFR/AFR× AFR/EAS;
E. 12 admixed cases with admixture ratio 50:25:25, where one parent is non-admixed, e.g. AFR/AFR ×EAS/EUR;
F. 12 admixed with admixture ratio 50:25:25, where both parents are admixed, e.g. AFR/EAS ×AFR/EUR;
G. 3 admixed cases with admixture ratio 25:25:25:25, e.g. AFR/EAS ×EUR/SAS;

For each of these 55 cases, we simulated 500 individuals in silico by picking four grandparents at random, creating mother and father from the grandparents, and creating a new individual from the parents, as described above.

### 2.3 Comparing results from admixture and recent-admixture

For a reference database from which we compute (or estimate) allele frequencies *p_mk_*, we can compute the estimate q from the admixture model as well as 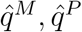 from the recent-admixture model, as described in (S5) and (S8). In order to compare the results from the admixture and recent-admixture model, we use deviations from the true *q* = (*q_k_*)_*k*=1,…, *K*_, where *q_k_* = 1 for a non-admixed individual from population *k, q_k_* = *q_k′_* = 0.5 for an admixed individual with parents from populations *k* and *k′*, etc. We use the Kullback-Leibler distance

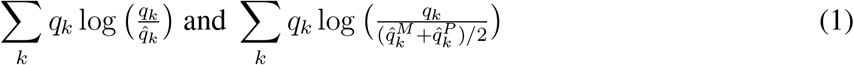

for the admixture model and the recent-admixture model, respectively. For estimating IA by using 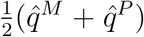, the recent-admixture model comes with the same number of model parameters as the admixture model. However, we stress that in the recent-admixture model, we obtain results for *q^M^* and *q^P^* separately, such that even more information than 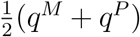 is contained in the estimates for this model.

### 2.4 Likelihood ratio test for recent admixture

We want to see if data *G* = (*G_m_*)_*m*=1…, *M*_ from a new trace fits significantly better to the recent-admixture model than to the admixture model. Since the admixture model is identical to the recent-admixture model for *q* = *q^M^* = *q^P^*, this amounts to a likelihood ratio test of *H*_0_: *q^M^* = *q^P^* against *H*_1_: *q^M^* ≠ *q^P^*. For this, we take the estimators 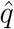 of *q* from iteration of (S5), and 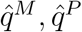 of *q^M^* and *q^P^* from iteration of (S8) and compute

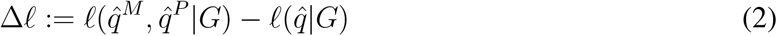

with *ℓ*(*q^M^, q^P^*|*G*) from (S6) and *ℓ*(*q*|*G*) from (S3). As usual in likelihood ratio tests, if Δ*ℓ* > *x* for some *x* (which needs to be specified), the recent-admixture model fits significantly better and we reject *H*_0_. If Δ*ℓ* ≤ *x*, we accept *H*_0_. Since markers are assumed to segregate independently, both 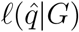 and 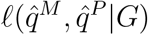 are sums over all M markers, such that we can even report the contribution of every AIM to Δ*ℓ*.

In an applied context, it is important to obtain a *p*-value from an observed value of Δ*ℓ*. The standard asymptotical theory for LR-tests (for large *M*) suggests that – under *H*_0_ – Δ*ℓ* is *χ*^2^-distributed with *K* – 1 degrees of freedom. However, in our applications, the number of markers is limited such that the asymptotic theory might not apply. Moreover, we see that the *χ*^2^-distribution turns out to be (i) too conservative in many cases and (ii) does not correctly account for the dependence of Δ*ℓ* on *q*; see Figure S9 in the SI. Therefore, we rely on simulations for obtaining a *p*-value in the following ideal way: First, we compute 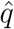 as well as 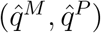. Then, we simulate individuals under *H*_0_, i.e. anciently admixed individuals with markers segregating independently and IA given by 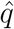, and compute the corresponding Δ*ℓ*-values for these individuals. Then, the *p*-value of the trace is the frequency of individuals with a higher Δ*ℓ* than the Δ*ℓ*-value of the trace. At least if 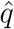 is close to *q*, this gives uniformly distributed *p*-values, which every statistical test aims for. However, we recognized that there are two factors confounding these reported *p*-values. First, in non-admixed traces, there is a chance that 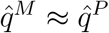 and small Δ*ℓ*, but still leading to a significant result because Δ*ℓ* is even smaller in most simulated samples. Since 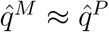 in these cases, recent admixture is not a useful conclusion, so we alter the procedure and report a *p*-value of 1 if 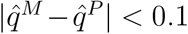. Second, in anciently admixed individuals originating from two or more populations, we overestimate the *p*-value by a factor at most two; see Section 3.4 below. Therefore, we alter the reported *p*-value and multiply the previously obtained value by two.

### 2.5 Data from the 1000 genomes project

In order to detect recent admixture in publicly available data, we downloaded 1000 Genomes data (phase 3) from ftp://ftp.1000genomes.ebi.ac.uk/vol1/ftp/release/20130502/, as well as information on the sampling locations from ftp://ftp.1000genomes.ebi.ac.uk/vol1/ftp/release/20130502/integrated_call_samples_v3.20130502.ALL.panel [22]. This is data from 661 individuals from Africa (AFR), 347 Admixed Americans (AMR), 504 East Asians (EAS), 503 Europeans (EUR) and 489 South-East Asians (SAS). The dataset comes with approximately 80 million SNPs. However, we use only a few of them known as the EUROFOR-GEN AIMset [23] and Kidd AIMset [24], respectively. The former comes with 128 SNPs, and we ignore seven tri-allelic SNPs (rsl7287498, rs2O69945, rs2l84O3O, rs433342, rs454OO55, rs5O3O24O, rsl24O2499), since our methods currently rely on bi-allelic SNPs. It was designed to distinguish Africa, Europe, East Asia, Native America, and Oceania, but was shown to perform well on the 1000 genomes dataset, also for distinguishing South Asia, even when ignoring the tri-allelic SNPs [7]. The latter comes with 55 bi-allelic SNPs and was introduced as a global AIMset differentiating between 73 populations. We note that this AIMset is part of the Verogen MiSeq FGx^™^ Forensic Genomics Solution.

The analysis of this dataset relies on allele frequencies used to estimate IA and PIA. Here, we use the samples of AFR, EAS, EUR and SAS. We did not use AMR since they are known to be admixed.

### 2.6 Analysis of two collected individuals with presumably recent admixture

Within a larger study about biogeographical inference, buccal swabs from two individuals were collected using a DNA-free swab (Sarstedt, Nümbrecht, Germany). The first individual reported one parent from Germany (Europe) and the other parent from the Philippines ((South-East)-Asian). The second individual reported a parent from Italy (Europe) and one from Venezuela (South America). Approval for collection and DNA analysis was obtained from the ethical committee of the University of Freiburg (414/18). DNA was extracted using the QIAamp Mini Kit (Qiagen, Hilden, Germany) and AIMs sequenced using the ForenSeq DNA Signature Prep Kit (Mix B) with the MiSeq FGx^®^ Reagent Micro Kit on a MiSeq FGx (all Verogen, San Diego, CA, USA). Sample preparation and sequencing was performed according to the Manufacturer’s recommendations. SNPs were analyzed and exported for inclusion in the model using the ForenSeq Universal Analysis Software (Verogen).

As a reference dataset for the analysis of the recent-admixture model (used for computing allele frequencies for continental populations), we use the Forensic *MPS AIMs Panel Reference Sets*, taken from http://mathgene.usc.es/snipper/illumina_55.xlsx which comes with the software Snipper [6]. This dataset contains data from the 1000 genomes project (504 out of 661 individuals from Africa (AFR) excluding the samples from African Caribbeans in Barbados and Americans of African Ancestry; 85 out of 347 Admixed Americans (AMR) only including Peruvians from Lima; 504 East Asians (EAS), 503 Europeans (EUR) and 489 South-East Asians (SAS)), as well as 13 Oceanian, Papua New Guinea, (OCE) samples from the Human Genome Diversity Panel. In the reference dataset, rs3811801 (contained in the ForenSeq DNA kit) is missing and therefore excluded from further analysis. This SNP has some discriminatory power for EAS (allele frequencies 1 (AFR); 1 (AMR); 0.49 (EAS); 0.99 (SAS)), as seen from the 1000 genomes data. Since data for rs1919550 and rs2024566 is missing for the Oceanic samples of the reference database, we also excluded these AIMs. Both only have low discriminatory power on a continental level. In total, this amounts to a total of 53 AIMs in the analysis, all of which are contained in the Kidd AIMset [24]. Allele frequencies are displayed in Figure S12.

### 2.7 Simulation of a three-islands model

We simulated genome-wide data from a sample, taken from a population genetic model with three islands, *A, B* and *C*, using [25]. Migration is such that only *A, B* and *B, C* are connected, but not *A, C*; see Figure 1(A). We used a migration rate of 10 diploid individuals per connected islands (in both directions) per generation. More precisely, we simulated a sample of 400 individuals per island, taken from a larger total population within each island, each with 20 recombining chromosomes, of which each carried about 2.5·10^4^ SNPs. From these ~ 5 · 10^5^ SNPs, representing part of the human genome, we selected 30 AIMs which are able to distinguish between *A* and *C*, i.e. with a large allele frequency differential; see Figure S1(A). From these 3 × 400 non-admixed individuals, we create first-generation admixed individuals in silico as described in Section 2.2 for the tests of performance of the AIMset.

**Figure 1:**
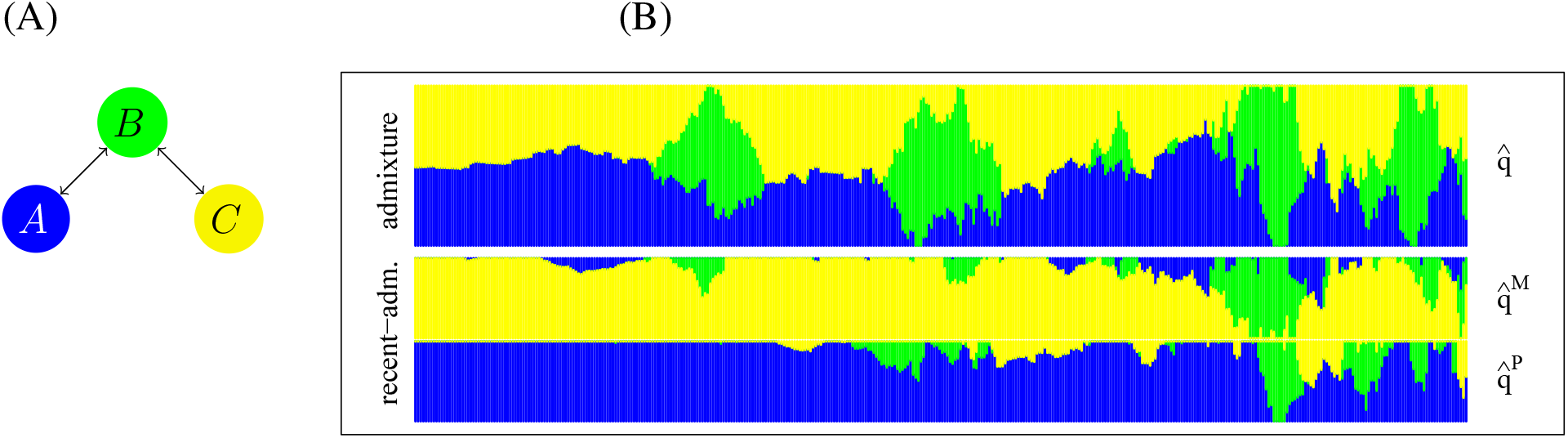
(A) Illustration of the population model for the simulations.Allele frequencies for the 10 AIMs used in the three island model. (B) In this barplot, every horizontal position gives one of 400 *A* × *C* first-generation-admixed individuals. In the top row, estimates 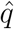 from the admixture model are given, and at the bottom, we give the corresponding estimates for 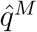 and 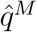 from the recentadmixture model.

## 3 Results

### 3.1 Analysis of a three-island model

We analyse the performance of the 30 AIMs with a large allele frequency differential between islands *A* and *C*. We find that these AIMs, when using a naive Bayes approach as in snipper [6], give a small (2%) misclassification error, even when adding individuals from B to the dataset; see Figure S1(B). However, for first-generation *A* × *C*-admixed individuals, the same classifier fails in many cases and gives population B as the best guess; see Figure S1(C).

We also use the admixture and recent-admixture model to estimate IA and PIA for both, non-admixed and first generation admixed individuals. We observe that the admixture model fails to give accurate estimates for IA in *A* × *C*-recently-admixed individuals for two reasons; see Figure 1(B). First, in almost all cases, admixture is not precise in estimating a 50% ancestry from *A* and *C*. Second, and more severely, in several individuals, IA wrongly predicts ancestry mostly in B, since allele frequencies in *B* are between *A* and *C*. Such wrong assignments occur less often when using recent-admixture and this model frequently correctly gives two parents of different ancestry, one mostly *A*, the other mostly *C*.

To get a picture of all non-admixed and recently admixed samples, we computed errors for estimating IA as given in (1) for the admixture and recent-admixture model. As described in the MM section, we average estimates 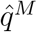 and 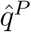 from the recent-admixture model, in order to compare to the true IAs. Figure S2 displays these errors in all cases including non-admixed individuals, and all three cases of recent-admixture. We note that on average, the recent-admixture model outperforms admixture on the *A* × *C* individuals, but gives less accurate estimates for the non-admixed *B*-individuals. In order to test if non-admixed *B*-individuals might be falsely classified as *A* × *C* individuals, we applied our statistical test for recent-admixture on these individuals. This resulted in a *p*-value below 0.05 for only 7 out of 400 non-admixed *B* individuals. To estimate the sensitivity of our test, we also applied it to 400 recently-admixed *A* × *C* individuals. Of these, 158 were correctly identified as recently admixed.

### 3.2 The recent-admixture model improves estimation accuracy of ancestry proportions for 1000 genomes project data

For comparing the accuracy of the admixture and recent-admixture model, we extended our analysis of the errors for estimating IA to the 1000 genomes dataset. We excluded all Admixed Americans (AMRs) since they are known to have an admixed background [2, 7] and do not form a well-defined own group. As the true IA, we use the continental origins as described in the dataset (AFR, EAS, EUR and SAS). This means e.g. that we set 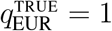 for a European sample in the dataset.

We ran three kinds of analyses. First, on the non-admixed samples. Second, we produced in silico recently admixed individuals with parents from the non-admixed samples (denoted AFR× EUR etc.) and ran the analysis on these samples. Third, the analysis was performed on second-generation admixed samples, i.e. grandparents were taken from the non-admixed samples (denoted AFR/EAS×EUR/SAS etc). In the first case, Figure S3 shows that the resulting errors for the admixture and recent-admixture model are almost identical, when using the EUROFORGEN AIMset. Overall, recent-admixture has a smaller error in 1843 out of 2157 cases, i.e. the hypothesis that the error for recent-admixture is at least as large as for admixture can be rejected (binomial test, *p* < 0.001). In the second case, Figure 2(A) shows clearly that errors for recent-admixture are smaller for all pairs of continents, when using the EUROFORGEN AIMset. More precisely, in 2262 out of 3000 individuals, recent-admixture is more accurate (*p* < 0.001). Third, for second-generation admixed individuals, Figure 2(B) displays errors in the cases (A)–(G) – recall from Section 2.2 – and shows that again, recent-admixture is more accurate, when using the EUROFORGEN AIMset. Here, recent-admixture outperforms admixture in 14881 out of 27500 cases (resulting in *p* < 0.001) A full list of 55 cases is displayed in Figure S4 in the SI. The corresponding results for the Kidd AIMset are similar and also found in the SI. We stress that the recent-admixture model not only gives significantly better estimates for IA, but also provides more information than the admixture model, since the genetic decomposition of both parents is estimated.

**Figure 2:**
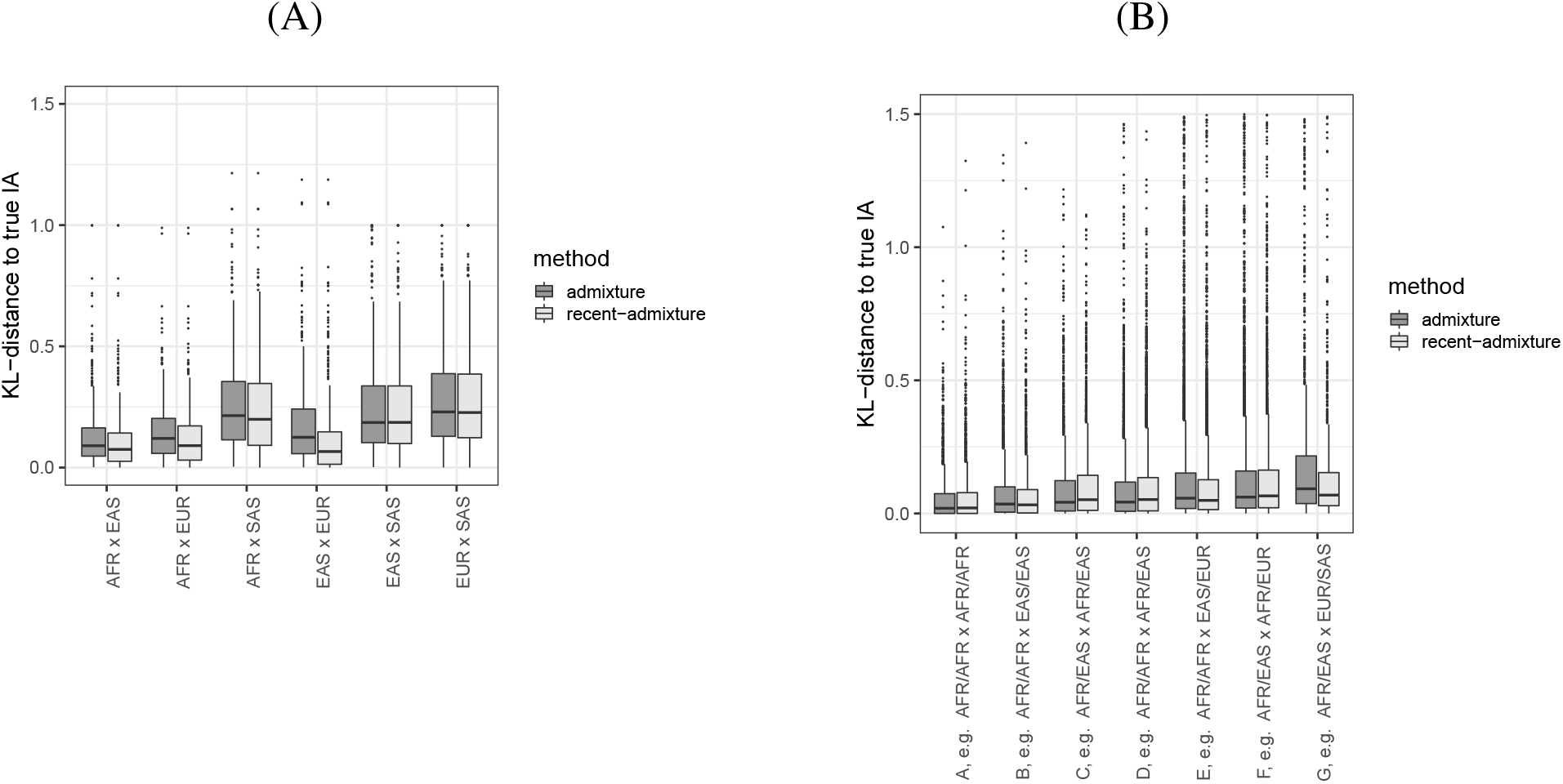
For all first generation admixed samples (A) and second generation admixed samples (B), we computed IA from the admixture 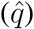 and recent-admixture 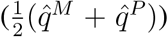 model using the EUROFORGEN AIMset. The distance to the true IA is computed as in (1). The cases in (B) are as described in Section 2.2.

### 3.3 Power of the Likelihood-ratio test for recent admixture

When fixing the minimal Δ*ℓ* for deciding if a sample is recently admixed in the likelihood-ratio test for recent admixture (as described in MM), we obtain the power of the test for all cases of recent admixture. For each case, we aim at distinguishing recent admixture with (*q^M^, q^P^*) from ancient admixture with *q* = (*q^M^* + *q^P^*)/2. Displaying the false positives (i.e. positively tested anciently admixed) against true positives (i.e. positively tested recently admixed) in all cases for all possible values of Δ*ℓ*, we obtain the Receiver-Operation-Characteristic (ROC) curve [26]. As we see in Figure 3, for the EUROFORGEN AIMset, the power of the test differs with the kind of recent admixture. The power is highest for first generation admixed (case (B)), and drops for one non-admixed parent (case (E)) and all grandparents from different continents (case (G)). It drops further if only half of the genome has two different ancestries (cases (D) and (F)). If the individual is not recently-admixed in first generation, but both parents are (case (C); note that this case is not recent-admixture, since *q^M^* = *q^P^* holds), or it is non-admixed (case (A)), recent admixture does not hold and the test has no power to detect these cases. The picture is nearly identical for the Kidd AIMset; see Figure S8.

**Figure 3:**
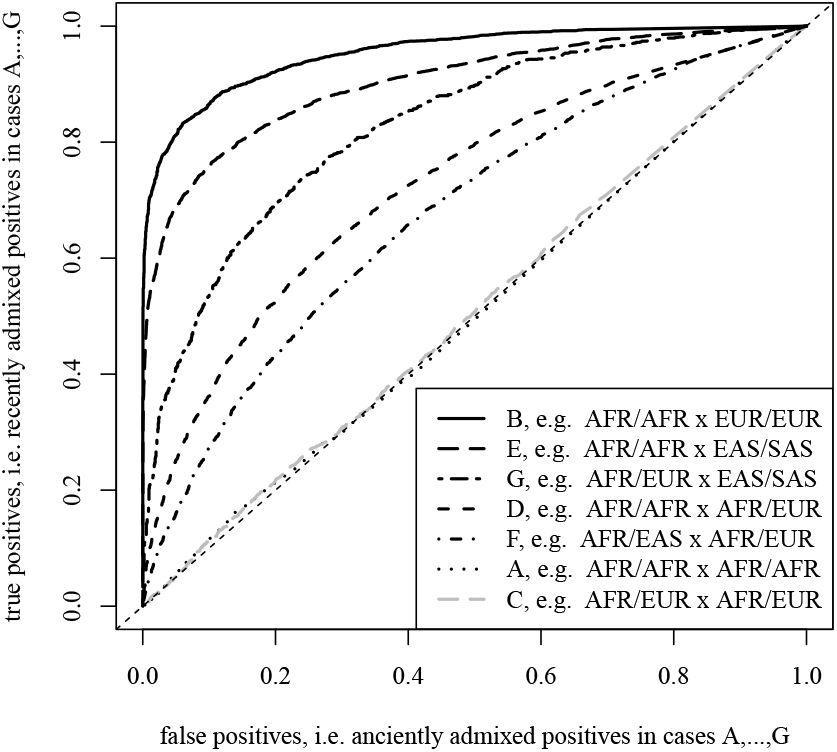
Using the EUROFORGEN AIMset, the ROC curve displays false positives (i.e. anciently admixed with the same *q*, against true positives (all cases of admixture in second generation)). In Figure 3 in the SI, a similar figure using the Kidd AIMset is given.

### 3.4 Simulating *p*-values

We checked if our approach of using simulations to study *p*-values is conservative. Both, the report of a *p*-value of 1 if |*q^M^* – *q^P^*| ≤ 0.1, and doubling the frequency of Δ*ℓ*-values above the Δ*ℓ*-value of the sample as reported *p*-value, are ad hoc. In order to analyse their performance, we use allele frequencies from the 1000 genomes dataset, again leave out AMR-individuals, and run simulations for individuals according to *H*_0_. So, for fixed *q*, we obtain anciently admixed genomes as described in Section 2.2, and run the computation of *p*-values. We investigate three cases, displayed in Figure S10 in the SI: (i) non-admixed individuals, where *q_k_* = 1 for some *k* ∈ {AFR, EAS, EUR, SAS}; (ii) anciently admixed individuals from two populations, where *q_k_q_k′_* = 0.5 for some *k, k′*, denoted AFR+EAS,…; (iii) q which is Dirichlet(1,…,1)-distributed (the uniform distribution on the simplex), denoted mixed. We see that *p*-values have a slight tendency to be not conservative only for samples mixed from all continents. All other cases, in particular, relevant cases of non-admixed individuals, have conservative *p*-values.

### 3.5 Structure as a confounding factor

In order to test if hidden structure in the reference populations can have confounding consequences for the LR test, we simulated a scenario with a strongly structured reference population. Therefore, we merged EAS and SAS in the 1000 genomes dataset into one reference population, but simulated non-admixed EAS, SAS individuals as well as other cases and applied the LR test. The simulated individuals from either EAS or SAS showed an increase in homozygous sites compared to the Hardy-Weinberg expectation from the structured reference. Therefore, hidden substructure in the reference populations causes the opposite effect than recent admixture would, which is an increase in heterozygous sites. Consequently, this scenario leads to conservative *p*-values, as can be seen from Figure S11 in the SI.

### 3.6 The LR test identifies recent admixture in the 1000 genomes dataset

From the 1000 genomes dataset, we highlight individuals which give significant results for the test of recent admixture for the EUROFORGEN and Kidd AIMsets. As trainingset, for estimating allele frequencies, we use the individuals from http://mathgene.usc.es/snipper/illumina_55.xlsx which are part of the 1000 genomes dataset; see Section 2.6. In Figure 4, we give the result of the most extreme individual (in the sense of the smallest *p*-value for both AIMsets observed in the whole sample), a male from the African American (ASW) population. We note that it is known that the ASW population is admixed [2], but until now, it has not been tested if admixture is recent. In Section S2.5 in the SI, we analyse all individuals from the dataset with a *p*-value below 5% for both, the EUROFORGEN and Kidd AIMset. In total, we find eight such individuals, four with African origin (all ASW) and four admixed Americans (two CLM, Colombian ancestry; one MXL, Mexican ancestry; one PUR, Puerto Rician ancestry). In Section S2.5 in the SI, we see that heterozygous sites are in fact the drivers of the large difference in log-likelihood between the recent-admixture and the admixture model.

**Figure 4:**
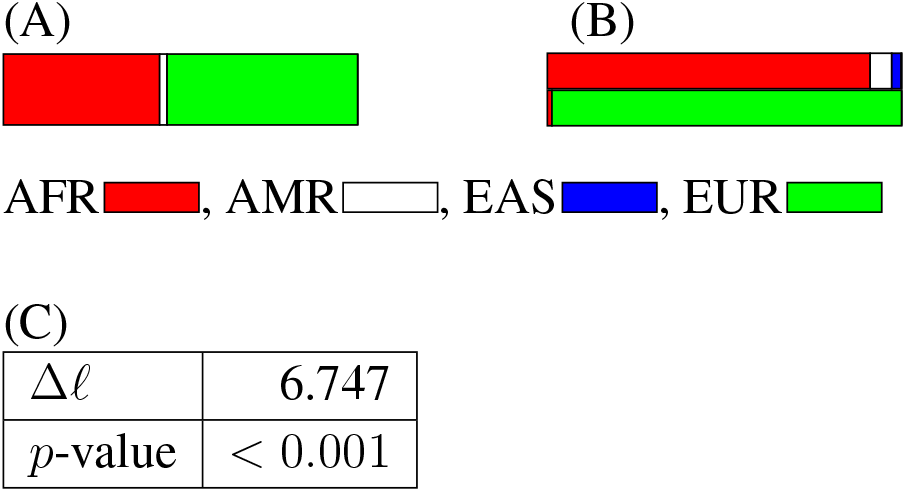
Analysis of NA20278 from the 1000 genomes dataset using the EUROFORGEN AIMset. (A) Estimated IA. (B) Estimated PIA. (C) *p*-value found by simulation, and difference in loglikelihood Δ*ℓ* from the likelihood-ratio test. More details on these contributions, broken down to single AIMs, is found in Section S2.5 in the SI.

### 3.7 Individuals with presumably recent admixture sequenced with the Verogen MiSeq FGx^™^ Forensic Genomics Solution

To further explore the performance of our method in an applied setting and to compare it to other current methods, we analyzed data from two individuals, generated with the Verogen MiSeq FGx^™^ Forensic Genomics Solution. For the German/Philippine female, when using a classification tool based on a naive Bayes approach (e.g. snipper), data from the 53 autosomal markers indicated a 61% chance to be European and 39% to be South-East-Asian (with reference samples from India, Pakistan etc). The ForenSeq Universal Analysis Software did not provide a clear classification result into one cluster of the training dataset, but the sample falls together with the Admixed American samples of the 1000 Genome project. The closest cluster is mainly comprised of samples from the 1000 genome populations from Puerto Rico and Colombia. The use of the admixture model leads to a mixed ancestry from Europe, East-Asian and Oceania; see Figure 5(A). Using the recent-admixture model, one parent with European and one parent with mainly an East-Asian ancestry are revealed which fits the self-declaration (Figure 5(B)). This individual has a *p*-value of 0.001, such that the recent-admixed model is favoured (Figure 5(C)).

**Figure 5:**
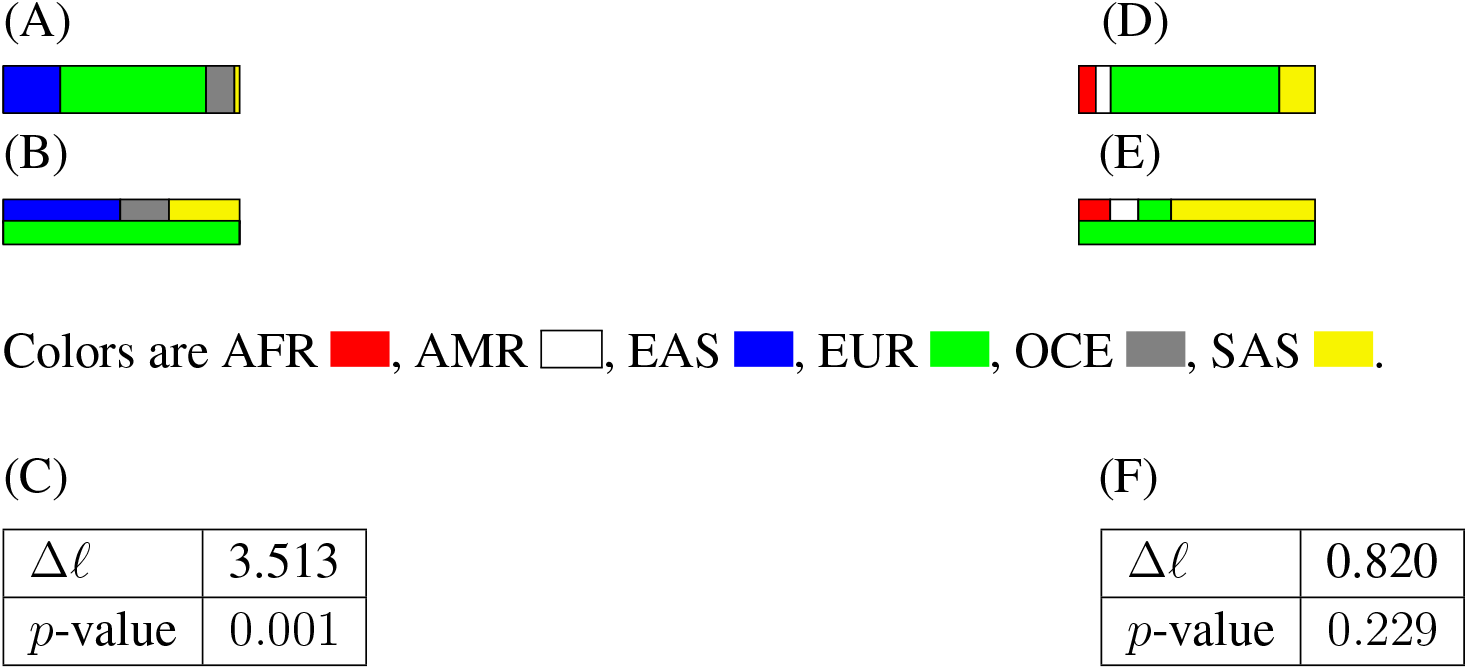
(A)–(C) Same as Figure 4, but for the first collected individual. (D)–(F) Same for the second collected individual. In Section S2.6 in the SI, contributions to Δ*ℓ* are broken down to single AIMs.

For the Italian/Venezuelan male, when using a naive Bayes classifier, his DNA is classified as European. The ForenSeq Universal Analysis Software provides a classification rather into the European cluster, but states the closest centroid in which also single reference samples from the 1000 genome project from European as well as Middle- and South-American ancestry fall into. The admixture model estimates mainly European ancestry, and contribution of South-East-Asian, and a small fraction African ancestry (Figure 5(D)). The recent-admixture model estimates one parent with European ancestry (explained by the Italian father) and one parent with mostly South-East Asian origin (Figure 5(E)). However, the corresponding *p*-value shows that this recent-admixture is not significant (Figure 5(F)).

### 3.8 Runtimes

The analysis of the admixture and recent-admixture model is fast. The main reason is that allele frequencies are only computed from a reference database (and not estimated on the fly, as in STRUCTURE and ADMIXTURE). As a consequence, runtimes scale linearly with the number of analysed traces. E.g. once allele frequencies for all AIMs from the reference dataset are given, the estimation of IA and PIA from one of the 500 · 55 = 27500 individuals in Figure 2, and which are created in silico, takes about 1.5 seconds per individuals on a standard laptop computer using the statistical language R. The most time-consuming procedure, however, is the computation of *p*-values via simulation. Here, for each simulated individual, again IA and PIA have to be computed, which then takes about three minutes per sample, if 100 individuals are simulated.

## 4 Discussion

Our first objective was to assess how recent admixture affects the outcome of standard methods of BGA inference. Recall that BGA is inferred either using a (all-or-nothing) classifier (as in SNIPPER [6]), or by estimating individual ancestry in a mixed membership model (as in STRUCTURE [4] and ADMIXTURE [3]). Using simulated island populations as well as data generated with the Verogen MiSeq FGx^™^ Forensic Genomics Solution, we provide in depth evidence that using classification software such as SNIPPER (all-or-nothing classifiers) is not suitable in recently admixed individuals, as has been suggested by [21]. Prestes et al. (2016) [27] notes that mixed membership models as used in STRUCTURE [4] and ADMIXTURE [3] are in many cases capable of inferring mixed ancestry in such individuals and [21] introduced a genetic distance algorithm, which improves results from ADMIXTURE in specific cases. However, we show that also the more sophisticated methods for BGA inference, STRUCTURE and ADMIXTURE, can be misleading in recently admixed individuals. These models fail because the assumption of equal chances for the two alleles at a locus to stem from any population is violated in recently admixed individuals. Here, as well as for various cases without recent admixture, the recent-admixture model on average gives more accurate results.

Our second objective was to improve the inference of BGA (or estimation of admixture composition, IA) and to develop a test to identify recently admixed individuals. We used the excess of heterozygote sites in recently admixed individuals – called the Wahlund principle [11] – in order to estimate parental admixture (see also [14]) and test for recent admixture. The recent-admixture model we present here is an extension of the admixture model. Whereas the admixture model assumes that an individual comes with its individual admixture (IA) from all reference populations, the extension presented here lies in the assumption that both parents can have different IA, which we call parental individual admixture (PIA). In particular, the recent-admixture model contains the well-established admixture model (see e.g. [4, 3]) as a special case. We note that the model cannot be easily generalized by e.g. introducing grand-parent individual admixture, since the signal of recent admixture of parents is lost on the level of the trace individual, at least if we assume linkage equilibrium between markers. Note that we still assume that AIMs are spread out in the genome, leading to independent segregation. This is in contrast to approaches using genome-wide dense SNP data, where linkage has to be taken into account [12, 13]. Another assumption made in our analysis is that allele frequencies in all populations are provided in a reference database. This is different from the approach taken in STRUCTURE, where IA and allele frequencies are estimated at the same time. In addition, we stress that the total number *K* of populations, which is a sensible parameter in various applications of STRUCTURE, see e.g. [28], here comes as a property of the reference database, i.e. is given externally. A consequence of the STRUCTURE approach is that the analyzed traces will shift the allele frequencies in the clusters that are posthoc assumed to represent the reference populations. We believe that in forensic genetics, the outcome of the analysis should not depend on e.g. the number of traces which are studied, and thus a method that relies on a reference data base should be preferred. A welcome side effect of leveraging a reference database is that our analysis is faster than STRUCTURE and ADMIXTURE. Since our method does not need genome-wide data, but only a few AIMs, analysis can be further accelerated and applied to forensic casework.

In inference of BGA in general, and for recent-admixture in particular, several hypothesis can be tested by a likelihood ratio (LR) test. For example, Tvedebrink and colleagues have recently developed statistical tests with the null hypothesis that the trace is a non-admixed sample from one of the populations in the reference database [16]. This test was extended in [17] to the null hypothesis that the trace is a recent admixture of the (non-admixed) parents from two populations in the reference database. In contrast, the LR test presented here tests if the recent-admixture model (with two parents with different IA) fits the data significantly better than the admixture model (where both alleles at a locus have the same chance given by IA to originate in any of the continents). As expected for a method that relies on a genome-wide increase in heterozygosity to detect recently admixed individuals, we see that heterozygote sites are responsible for the biggest contribution to the increase in likelihood in the recent-admixture model. Related to this Wahlund effect are thoughts on confounding factors for inference of recent-admixture. At least, we have shown that hidden structure within the reference database and the trace does not confound recent-admixture, since hidden structure leads to an excess of homozygotes (rather than an excess of heterozygotes). At least, only few mechanisms are known to inflate the number heterozygotes. As for heterozygote advantage, it is hardly conceivable that such a locally (in the genome) acting selective force confounds genome-wide patterns of recent admixture

For determining *p*-values and the specificity of the LR test, we rely on simulations. Thus, the asymptotic LR-theory does not apply and the distribution of the LR-test statistics depends on the true IA. On the way, we had to make some ad-hoc decision rules, one of them being that the estimated IA for the two parents has to be sufficiently different. Here, a better understanding of the theoretical distribution of the LR test statistics, would certainly be desirable. The sensitivity or power of our LR test to detect recent-admixture is highest if the two parents have non-admixed IA from different populations. Other cases, such as four grand-parents coming from four different populations, unsurprisingly give a lower power.

In our real-world samples, we detected signs of recent-admixture both in the 1000 genomes dataset and the collected individuals. Note that the 1000 genomes project specifically targeted individuals that are most likely to have all of their grandparents in only one location or population [29]. Nevertheless, we find striking evidence for recent admixture in some African individuals from the southwest of America, as well as for some admixed Americans. For the two presumably admixed samples that were analyzed with ForenSeq, the analysis of recent-admixture was compared with the self-reporting of the individuals. The ancestry information is therefore based on the assumption of having a biologically correct family tree and correct information on birth place and ancestry. In case of the female individual, Philippine and German ancestry were declared for the last four generations. This individual was successfully identified as recently admixed. The other individual, with self-reported maternal Venezuelian ancestry of the last two generations, was “insecure” about the generation before. Both IA and PIA suggested a proportion of SAS ancestry for this individual, which seems misleading. However, it has been noted that admixed South American populations have a high genetic overlap with SAS populations, making it difficult to differentiate these populations even when analyzed with genome-wide, dense SNP data [30]. This suggests that the AIM based approaches that we applied might not be able to resolve these differences and misidentify the South American ancestry as SAS ancestry in this sample. In addition, the AIMset used here is not perfect in distinguishing EUR and SAS, suggesting that an extended AIMset could lead to a significant result for recent-admixture. Morevoer, we have no population from Venezuela in the reference database which might have added to the apparent misassignment of both, IA and our newly developed PIA. Because Europeans contributed substantially to the genetic make up of the Venezuelan population via ancient admixture [31], our method might lack power to detect the recent admixture event in this case. At least, the recent-admixture model correctly picks up these uncertainties of the reference database, and finally leads to a non-significant result for recent-admixture.

## Acknowledgements

We thank Denise Syndercombe Court for comments on an earlier version of the manuscript, and two anonymous reviewers for several suggestions, in particular for pushing us to report *p*-values. The sequencing results are part of a larger study which is partly funded by the Wissenschaftliche Gesellschaft Freiburg.

## Supporting Information

### S1 Theory

We write down the admixture model and derive a method to estimate Individual Admixture (IA) in the case when allele frequencies within populations are not updated. In addition, we give the recent-admixture model, where an individual is allowed to have parents with different admixture proportions. We take the following notation for the reference database:

*K*: number of ancestral populations,
*M*: number of markers,
*p_mk_*: frequency of allele 1 at (bi-allelic) marker m in population *k*.

In addition, we consider one additional diploid genome (called the trace) (*G*_*m*1_, *G*_*m*2_)_*m*=1,…, *M*_, or (*G_m_*)_*m*=1,…, *M*_ with *G_m_* = *G*_*m*1_ + *G*_*m*2_ if phase is not known. The goal is to estimate admixture proportions (*q_k_*)_*k*=1,…, *K*_ (or 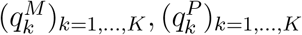) of this additional genome. Recall that we take the approach from [1] and [2] and do not update *p_mk_*’s during the analysis.

#### S1.1 The admixture model

In [3], [4], [5] and elsewhere, the main goal is to maximize the log-likelihood (see also (1) and (2) of [5])

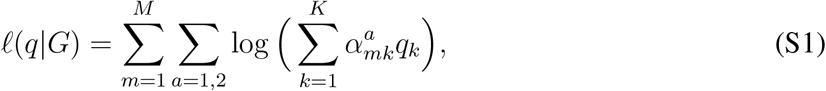

where

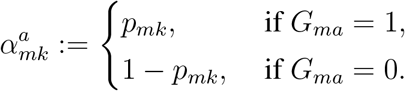

is the frequency of the observed allele in copy *a* of marker m in population *k*. (Note that *e*^*ℓ*(*q*|*G*)^ is the probabiltiy of observing (*G_ma_*)_*m*=1,…, *M*;*a*=1,2_, if every allele is picked independently from population *k* with probability *q_k_*.) Assuming that phase is not known, and with

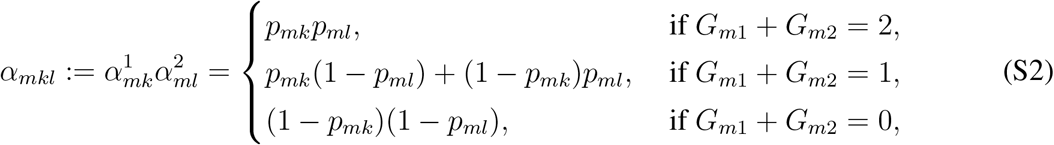

note that the log-likelihood can as well be written as

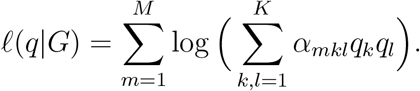

We set 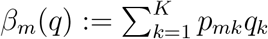 and analyse the last sum by distinguishing the case *G_m_* = 2, where

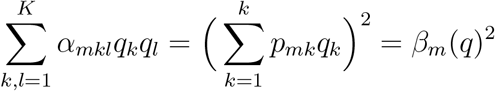

while for *G_m_* =1 we find

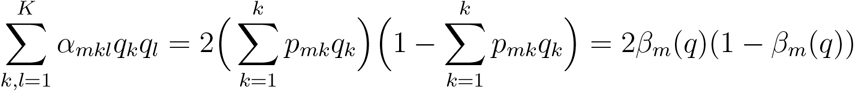

and for *G_m_* = 0

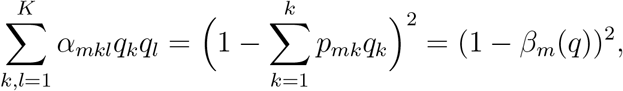

such that

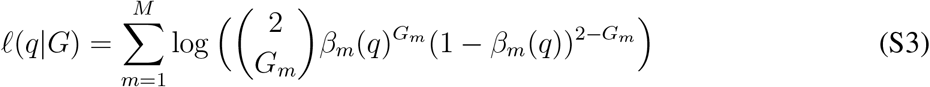

##### Lemma S1.1.

*The maximum of q* ↦ *ℓ*(*q*|*G*) *under the constraint* 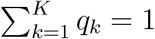 *solves*

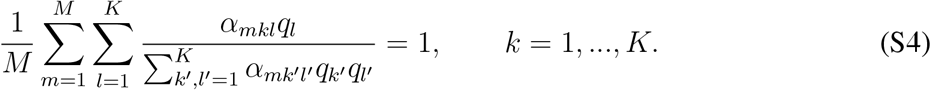

##### Remark S1.1.

1. Assume we add a set of *Ancestry Uninformative Markers*, i.e. a set of markers 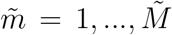, with frequencies not depending on the population, i.e. 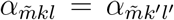 for *k, l, k, l″* = 1,…, *K*. For these markers,

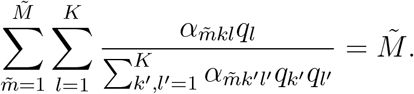 This implies that *q* is a solution of (S4) without these markers iff *q* is a solution of (S4) if these markers are included. This might be reassuring.
2. Let us have a closer look at the left hand side of (S4). We set 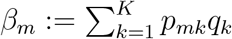. For *G_m_* = 2,

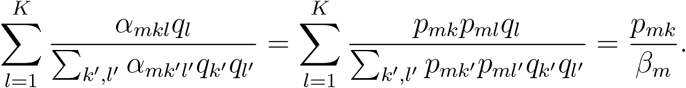 For *G_m_* = 1, we have

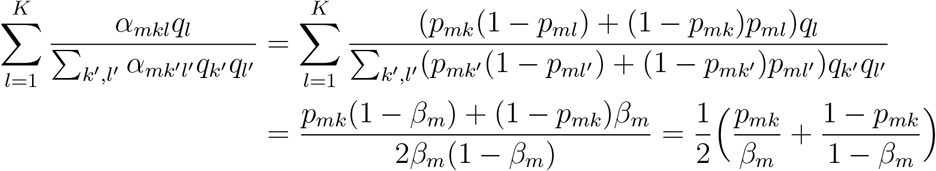

and for *G_m_* = 0

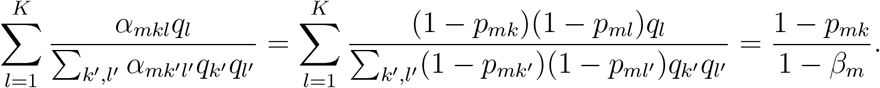 In total, this gives that *q* needs to solve

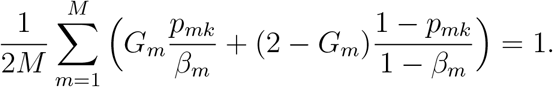 In order to find a solution, we reformulate as the fixed point equation

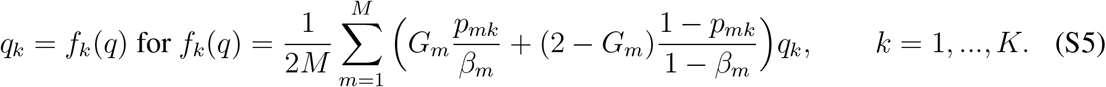 This can be done numerically by iterating *q*_*n*+1_ = (*f_k_*(*q_n_*))_*k*=1,…, *K*_ until convergence. (Note, however, that we do not have a formal proof of convergence for this iteration. At least, in our numerical experiments, convergence – in the sense that we continue the iteration until |*q*_*n*+1_ – *q_n_*| < 10^-6^ – always happened.) We note that this approach is essentially the same as in the EM-algorithm from [5], but combining the expectation and maximization steps, since we do not update allele frequencies. In addition, although maximizing (S3) could also be handled using a Newton method as in [3], this approach has the advantage that *q_n_*’s are positive in all steps, and the sum of all entries in *q_n_* is always 1. Moreover, the iteration is computationally fast if only a small or moderate number of markers is considered.

*Proof of Lemma S1.1*. We use the theory of Lagrange multipliers, since we need to maximize *ℓ* over q under the constraint 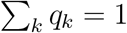. Since

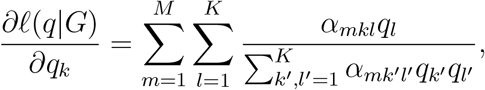

we have to solve the system of equations

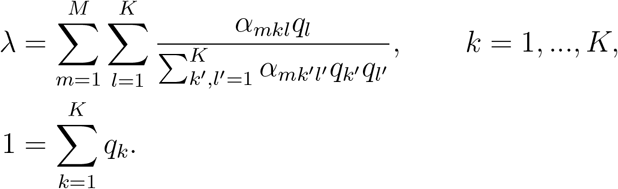

It is easy to eliminate *λ*, since the last equation gives

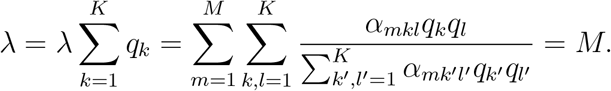

So, we are left with finding *q* = (*q_k_*)_*k*=1,…, *K*_ such that

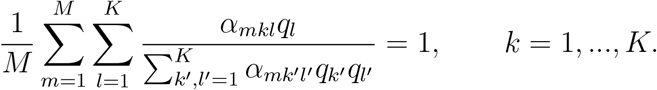

#### S1.2 The recent-admixture model

For the recent-admixture-version, we want to estimate 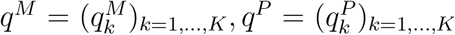, where 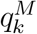 and 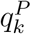 are the fractions of the genomes of the mother and father, respectively, which come from population *k*. This assumption implies that the log-likelihood is

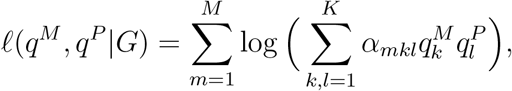

where *α_mkl_* is given as in (S2). As is apparent from the log-Likelihood function, the recent-admixture model generalizes the admixture model. Put differently, choosing *q^M^* = *q^P^* = *q* in the recent-admixture model gives the admixture model. We note that again, the log-likelihood can be written differently,

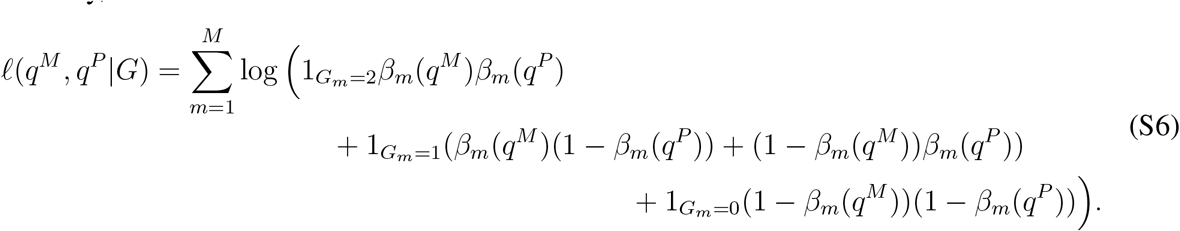

##### Lemma S1.2.

*The maximum of* (*q^M^, q^P^*) ↦ *ℓ*(*q^M^, q^P^, G*) *under the constraint* 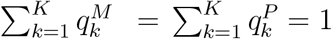 *solves*

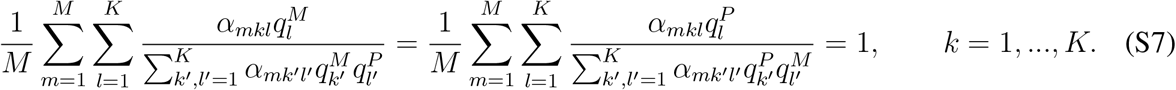

##### Remark S1.2.

1. Note that (S7) is symmetric in *q^M^* and *q^P^*, i.e. if (*q^M^, q^P^*) solve (S7), another solution is given by (*q^P^, q^M^*).
2. Again, we can turn (S7) into fixed point equations. To derive it, we again have a closer look at the left hand side of (S7). For 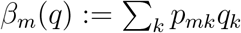, we have for *G_m_* = 2

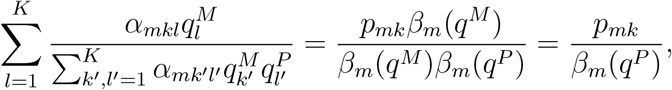

for *G_m_* = 1

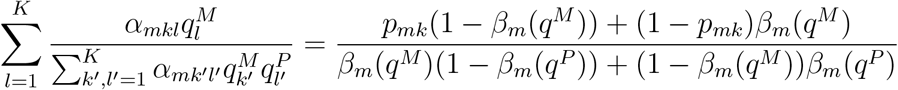

and for *G_m_* = 0

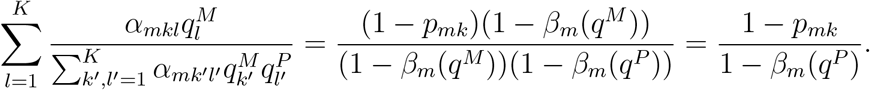 So, we suggest to iteratively compute

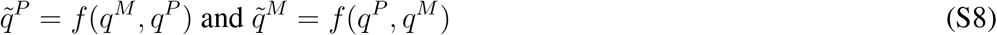

for *f*(*q, q′*) = (*f_k_*(*q, q′*))_*k*=1,…, *K*_ with

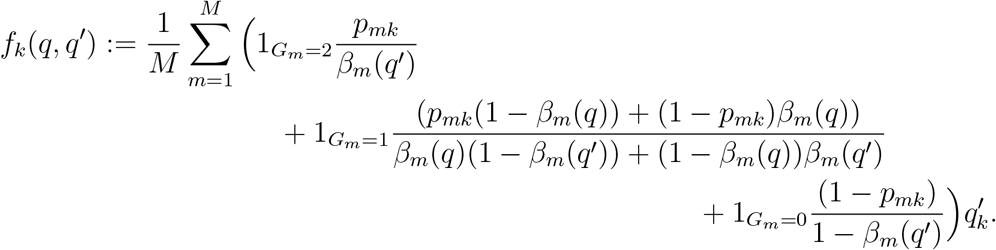

*Proof of Lemma S1.2*. Again, we use Lagrange multipliers. Since

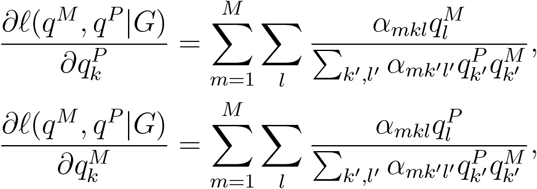

we have to solve the system of equations

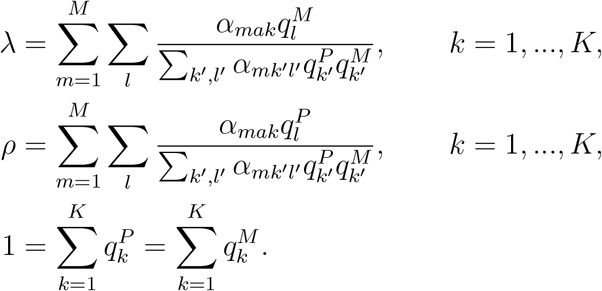

It is easy to eliminate *λ* and *ρ*, since

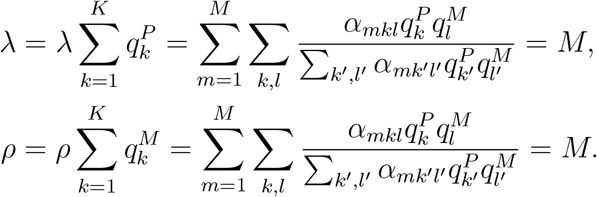

So, we are left with finding *q^P^* and *q^M^* such that

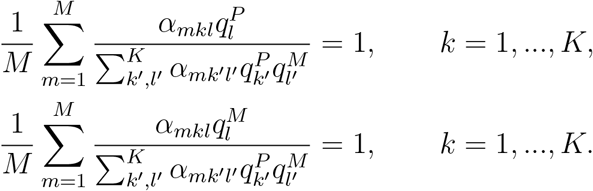

### S2 Additional results

#### S2.1 Some showcases from simulations

**Figure S1:**
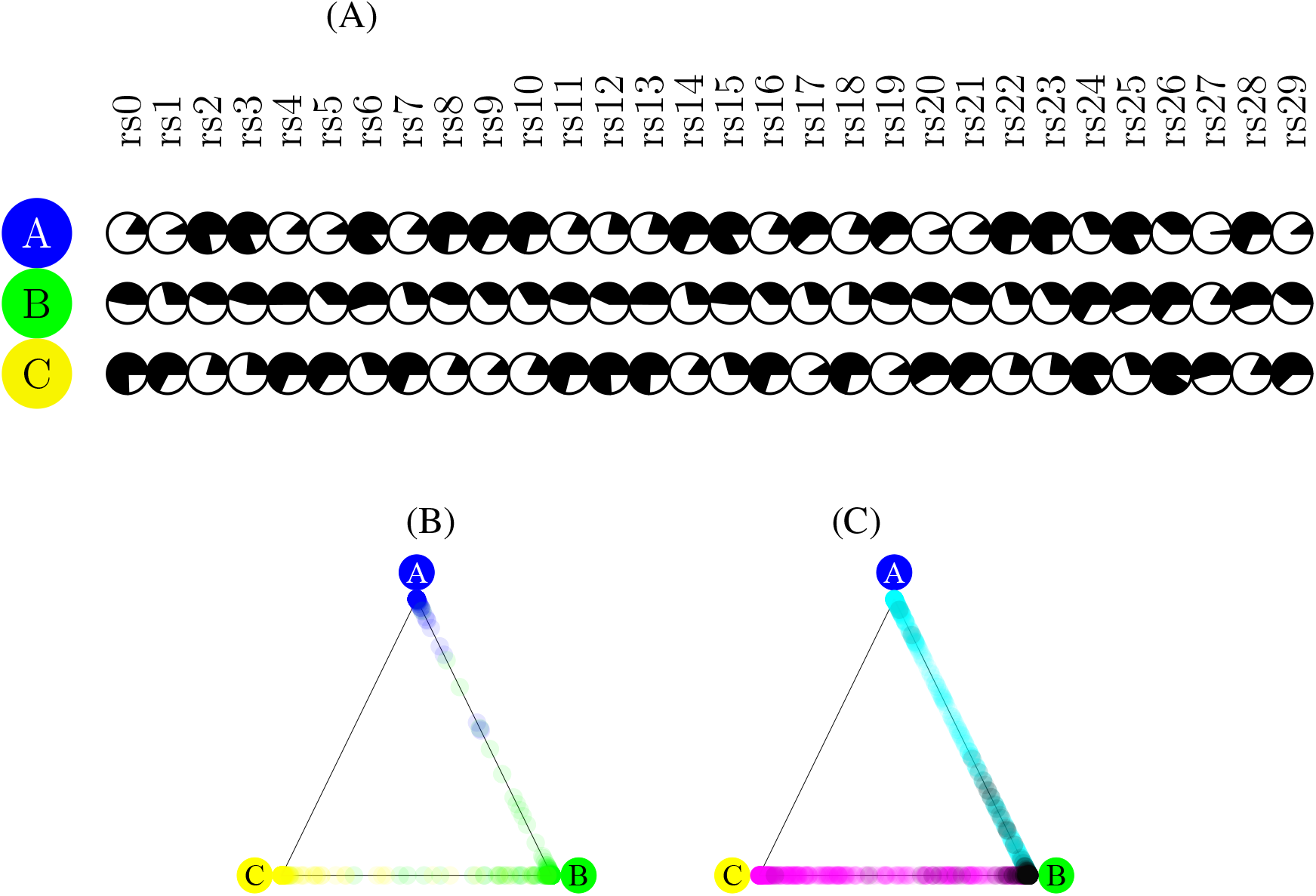
(A) From the simulations, we picked the 30 AIMs with the largest allele frequency differential between *A* and *C*. (B) When using all-or-nothing classifiers such as Snipper [6] based on the allele frequencies from (A), the probabilities of belonging to population *A, B* and *C* are plotted for non-admixed individuals. (C) Same as in (B) for first-generation admixed individuals from *A* × *B* (light black), *A* × *C* (black) and *B* × *C* (pink). Remarkably, all *A* × *C* are wrongly classified into *B*.

**Figure S2:**
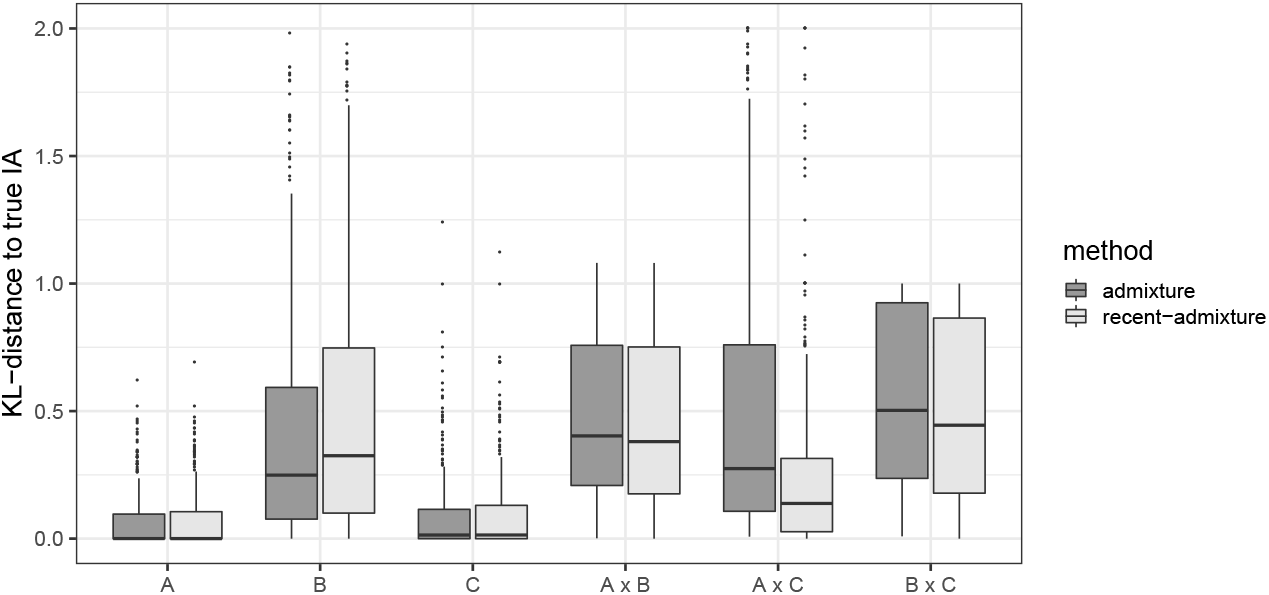
To get a picture of all non-admixed and recently admixed samples, we computed errors for estimating IA as given in (1) for the admixture and recent-admixture model. As described in the MM section, we average estimates 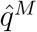 and 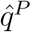 from the recent-admixture model, in order to compare to the true IAs.

#### S2.2 Estimation accuracy

For non-admixed and first generation admixed individuals, all cases (parents come from different continents) are described in the main text. For second generation admixed individuals, we have several cases, depending on the origin of the grand-parents. When data from four populations (AFR, EAS, EUR, SAS; see below) is available, we have the following cases:

A. 4 non-admixed cases with IA 100:0: AFR/AFR×AFR/AFR, EAS/EAS×EAS/EAS, EUR/EUR× EUR/EUR, SAS/SAS×SAS/SAS;
B. 6 admixed cases with IA 50:50, where both parents are non-admixed: AFR/AFR× EAS/EAS, AFR/AFR× EUR/EUR, AFR/AFR× SAS/SAS, EAS/EAS× EUR/EUR, EAS/EAS× SAS/SAS, EUR/EUR× SAS/SAS;
C. 6 admixed cases with IA 50:50, where both parents are admixed: AFR/EAS × AFR/EAS, AFR/EUR × AFR/EUR, AFR/SAS × AFR/SAS, EAS/EUR × EAS/EUR, EAS/SAS×EAS/SAS, EUR/SAS×EUR/SAS;
D. 12 admixed cases with IA 75:25: AFR/AFR× AFR/EAS, AFR/AFR× AFR/EUR, AFR/AFR× AFR/SAS, EAS/EAS× EAS/AFR, EAS/EAS× EAS/EUR, EAS/EAS×EAS/SAS, EUR/EUR× EUR/AFR, EUR/EUR ×EUR/EAS, EUR/EUR× EUR/SAS, SAS/SAS× SAS/AFR, SAS/SAS× SAS/EAS, SAS/SAS× SAS/EUR;
E. 12 second generation admixed with IA 50:25:25, where one parent is non-admixed: AFR/AFR× EAS/EUR, AFR/AFR× EAS/SAS, AFR/AFR× EUR/SAS, EAS/EAS× AFR/EUR, EAS/EAS×AFR/SAS, EAS/EAS×EUR/SAS, EUR/EUR×AFR/EAS, EUR/EUR×AFR/SAS, EUR/EUR×EAS/SAS, SAS/SAS×AFR/EAS, SAS/SAS×AFR/EUR, SAS/SAS×EAS/EUR;
F. 12 second generation admixed with IA 50:25:25, where both parents are admixed: AFR/EAS ×AFR/EUR, AFR/EAS × AFR/SAS, AFR/EUR× AFR/SAS, EAS/AFR×EAS/EUR, EAS/AFR×EAS/SAS, EAS/EUR×EAS/SAS, EUR/AFR×EUR/EAS, EUR/AFR×EUR/SAS, EUR/EAS×EUR/SAS, SAS/AFR×SAS/EAS, SAS/AFR×SAS/EUR, SAS/EAS × SAS/EUR;
G. 3 second generation admixed with IA 25:25:25:25: AFR/EAS×EUR/SAS, AFR/EUR×EAS/SAS, AFR/SAS×EAS/EUR;

As can be seen in Figure S4, the recent-admixture model on average does not give worse estimates than the admixture model, and outperforms the admixture-model in several cases; see also Figure 2 in the main text. For the Kidd AIMset, see Figures S6 and S7.

##### EUROFORGEN AIMset

**Figure S3:**
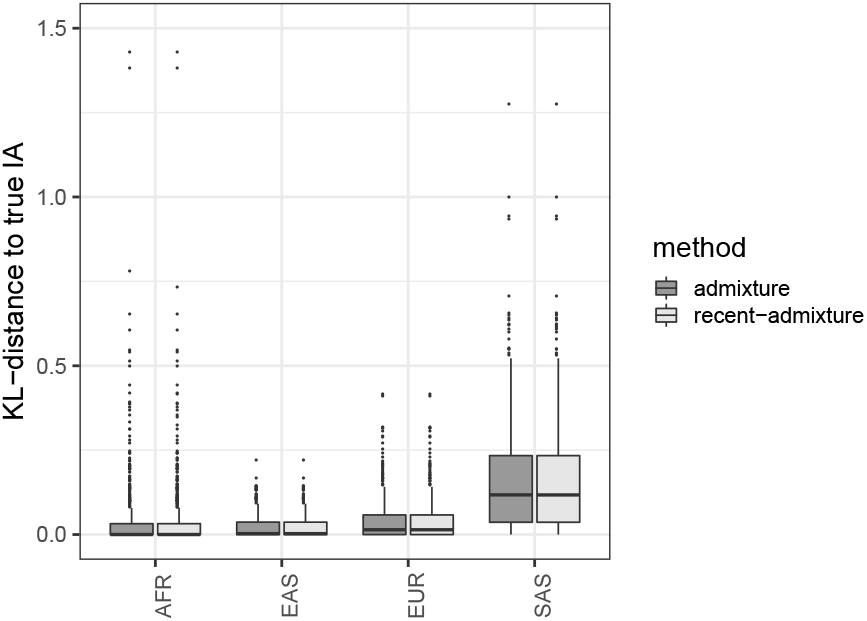
For all non-admixed samples from the 1000 genomes dataset, we computed IA from the admixture and recent-admixture model. The distance to the true IA is computed as in (1).

**Figure S4:**
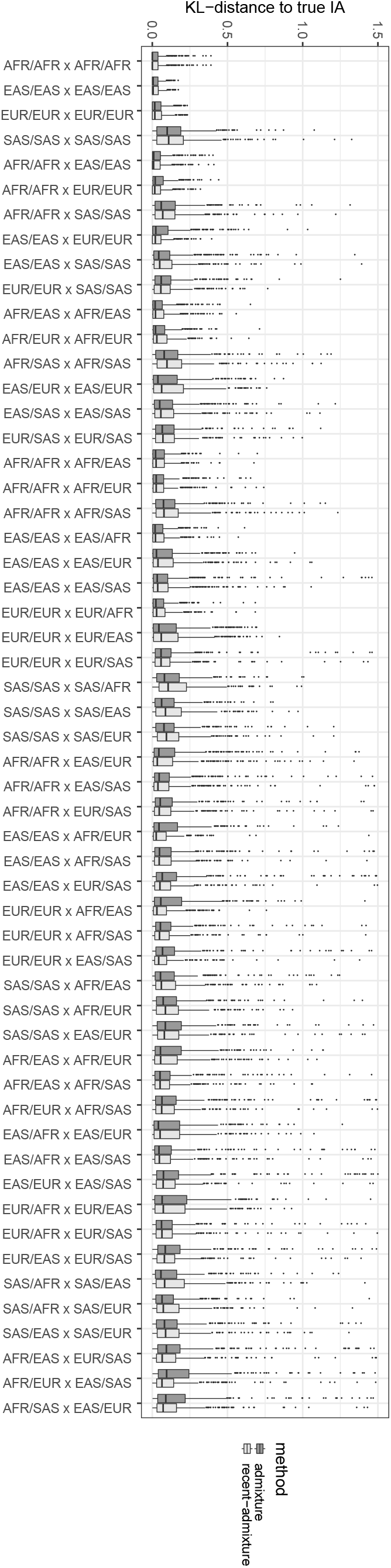
For all cases of second generation admixed individuals, we compare estimation accuracy of IA using the EUROFORGEN AIMset.

##### Kidd AIMset

For the Kidd AIMset, recent-admixture produces an error in the non-admixed samples (see also Figure S5), which is smaller than the error for the admixture-model in 1394 out of 2157 cases, so the hypothesis that the error is worse for recent-admixture can be rejected (*p* < 0.001). For first order admixed samples, 2381 out of 3000 cases have a smaller error under recent-admixture (which gives *p* < 0.001; see also Figure S6.A. Last, for second-generation admixed individuals, recent-admixture is more accurate in 16116 out of 27500 cases (*p* < 0.001); see also Figures S6.B and S7.

**Figure S5:**
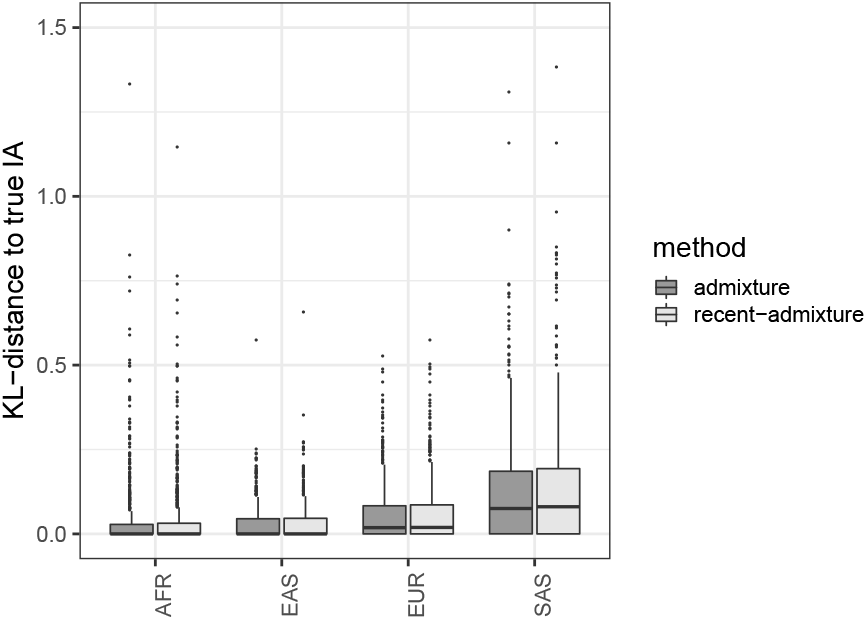
Same as in Figure S3, but for the Kidd AIMset.

**Figure S6:**
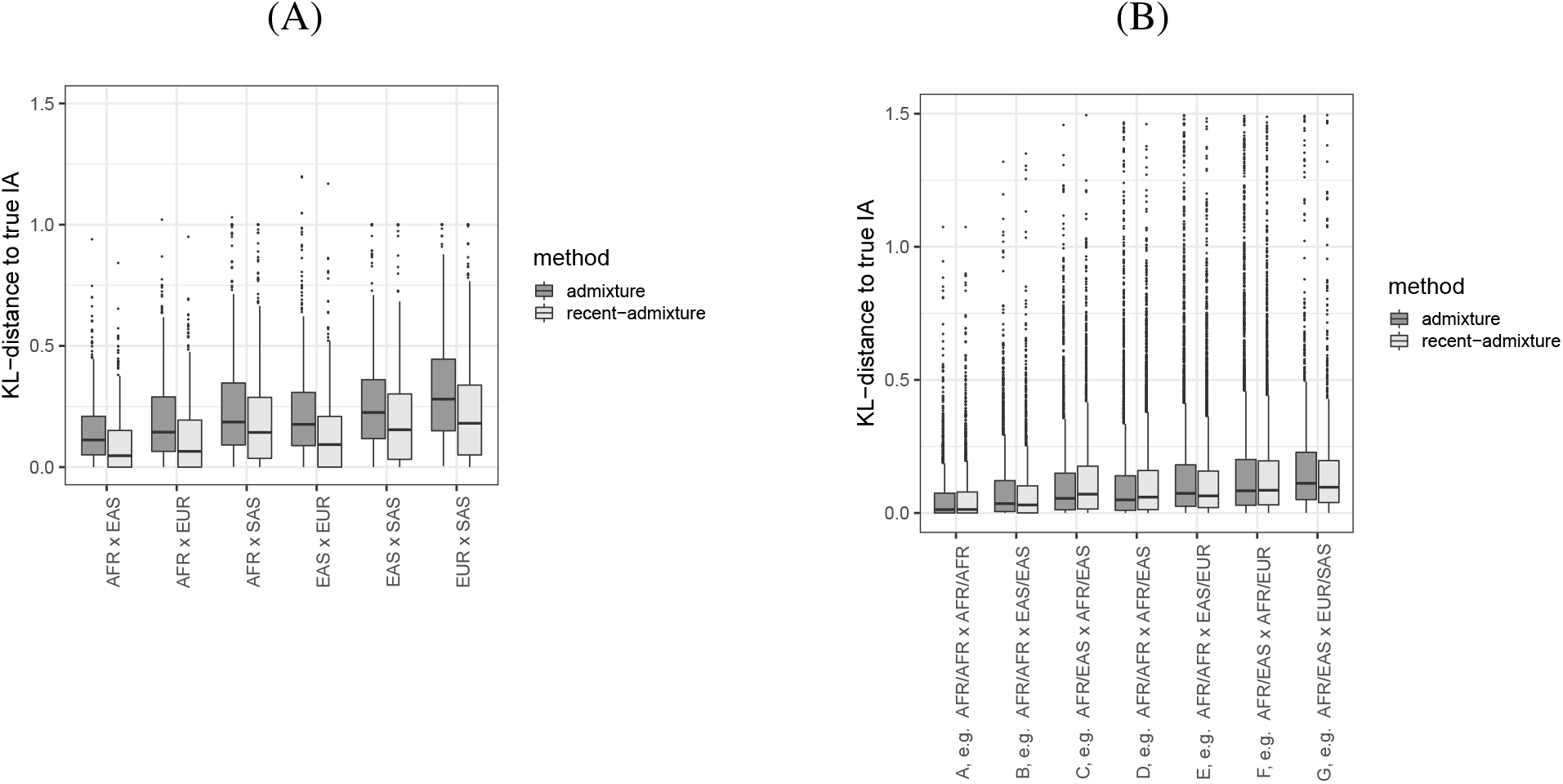
Same as in Figure 2, but using the Kidd AIMset.

**Figure S7:**
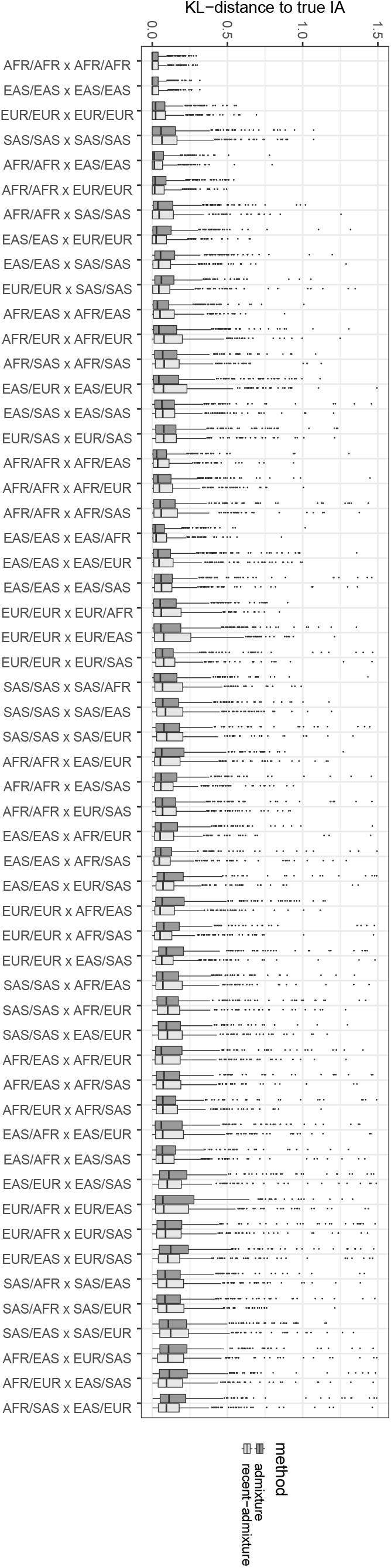
Same as in Figure S4, but using the Kidd AIMset.

**Figure S8:**
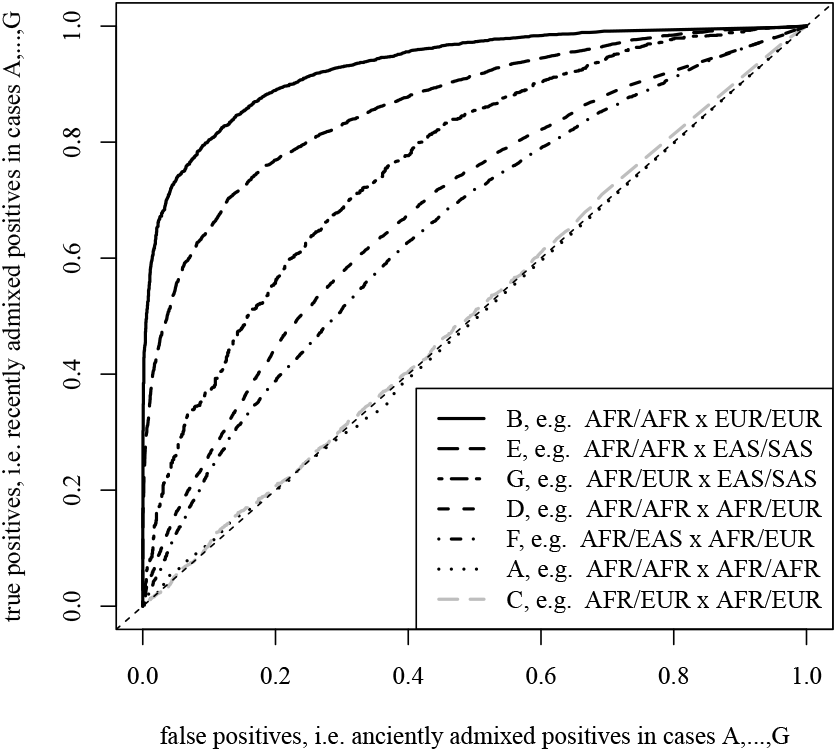
Same as in Figure 3, but using the Kidd AIMset.

### S2.3 Simulating *p*-values and hidden structure

**Figure S9:**
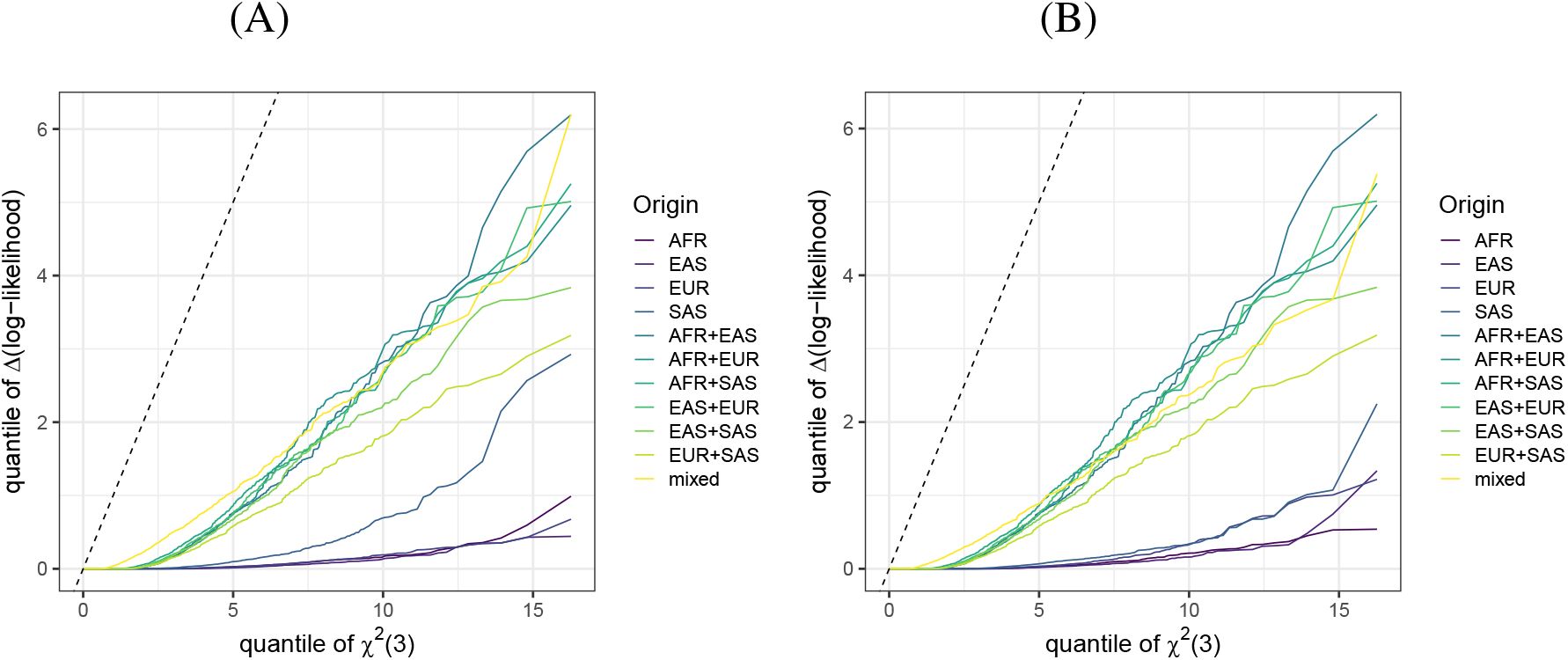
Quantile-quantile plot of Δ*ℓ* against a *χ*^2^ distribution with three degrees of freedom. AFR,… are simulated under *q*_AFR_ = 1,… from the allele frequencies. AFR+EAS,… are simulated under *q*_AFR_ = *q*_EAS_ = 0.5,… For Dirichlet-distributed *q*, we have the mixed case. The distribution of Δ*ℓ* not only is far from *χ*^2^ (3)-distributed, but also depends on *q*. (A) For the EUROFORGEN AIMset with classes AFR, EAS, EUR and SAS; (B) For the Kidd AIMset.

**Figure S10:**
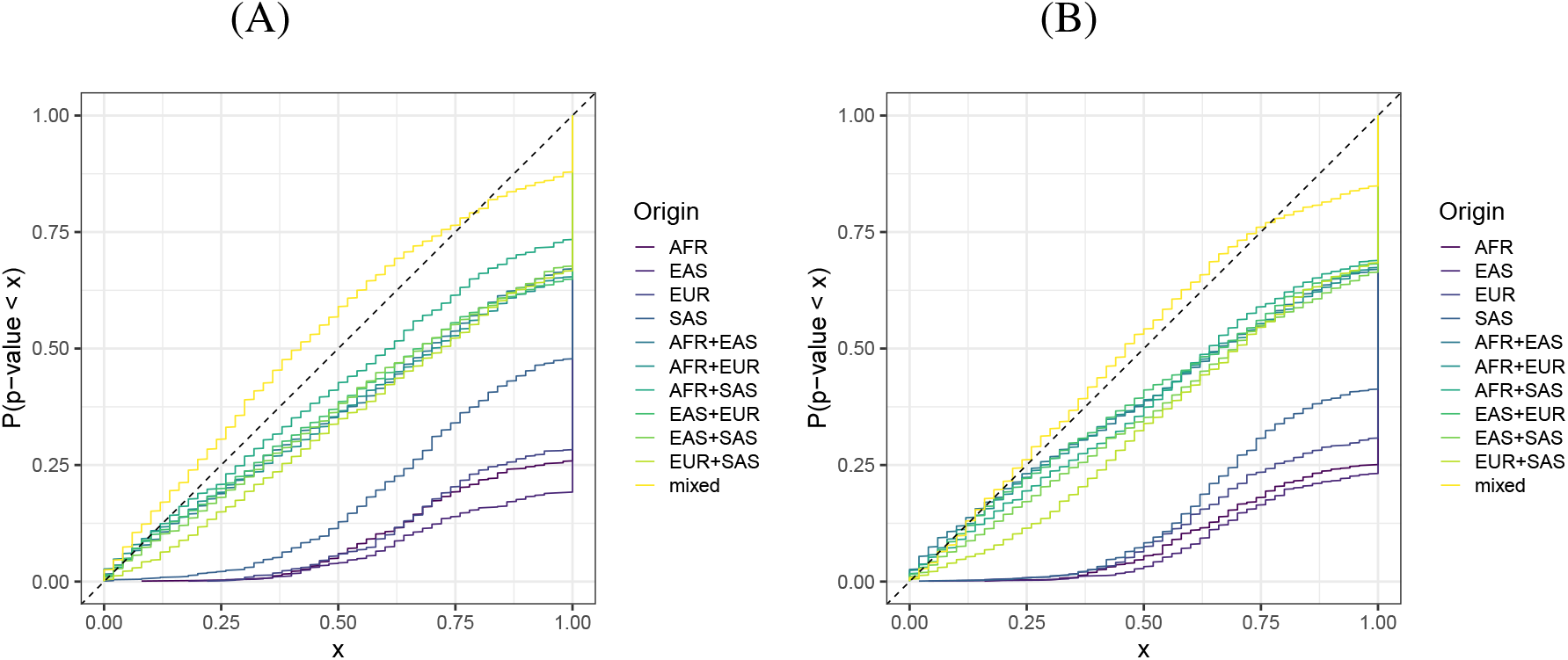
Distribution of *p*-values for several types of individuals. Data used is as in Figure S9. If the curve is below *f*(*x*) = *x*, we are facing a conservative test, since there is at most a fraction *x* of *p*-values below *x*. (A) For the EUROFORGEN AIMset with classes AFR, EAS, EUR and SAS; (B) For the Kidd AIMset.

**Figure S11:**
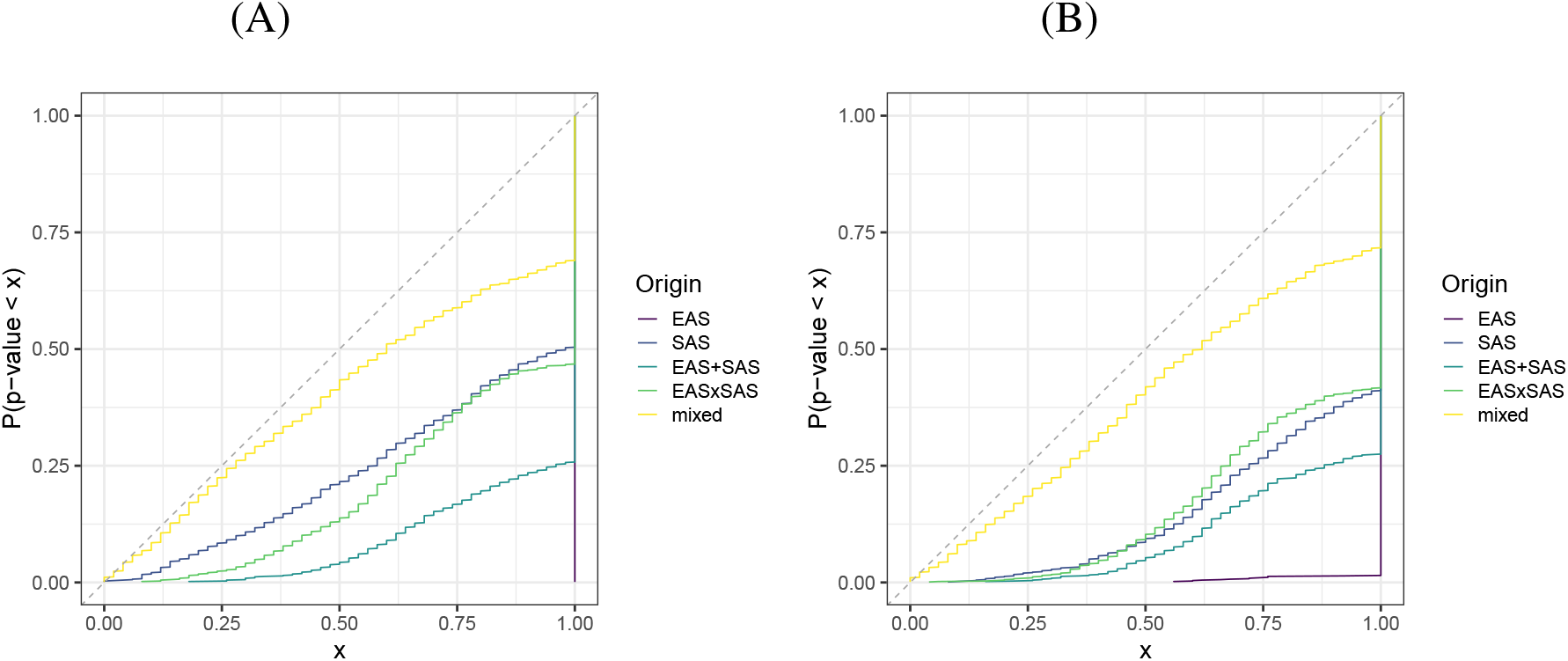
Same as in Figure S10, but EAS and SAS was combined in the reference database, i.e. it has hidden structure. (A) For the EUROFORGEN AIMset. (B) For the Kidd AIMset.

### S2.4 More details for the AIMs analysis of two collected individuals with recent admixture

**Figure S12:**
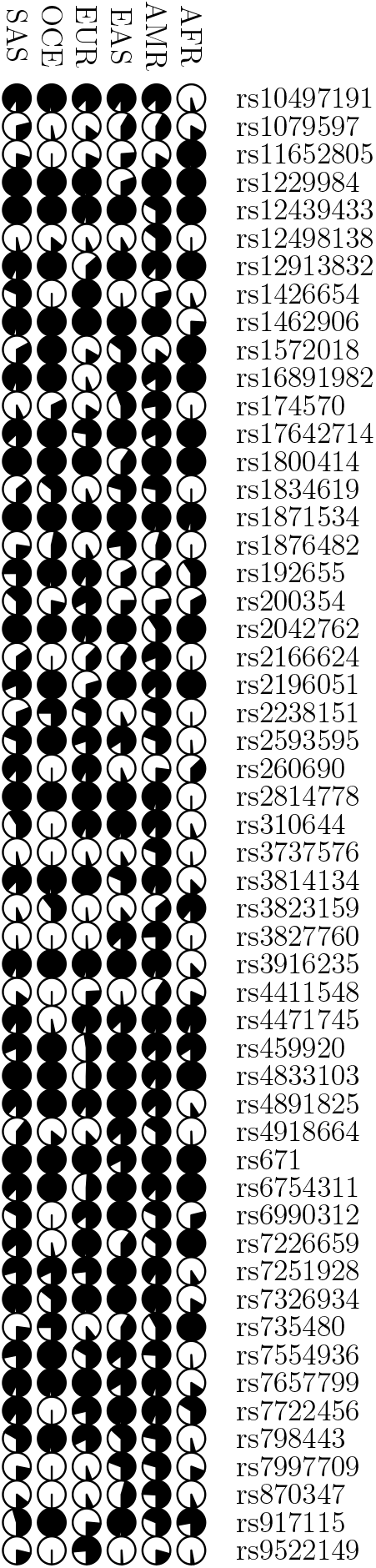
Allele frequencies for six populations for all 53 AIMs from the Forensic *MPS AIMs Panel Reference Sets*, taken from http://mathgene.usc.es/snipper/illumina_55.xlsx, which are used in the analysis of the collected individuals.

### S2.5 Recent admixture in the 1000 genomes dataset

#### Analysis of HG01092, (AMR, PUR)

##### EUROFORGEN AIMset

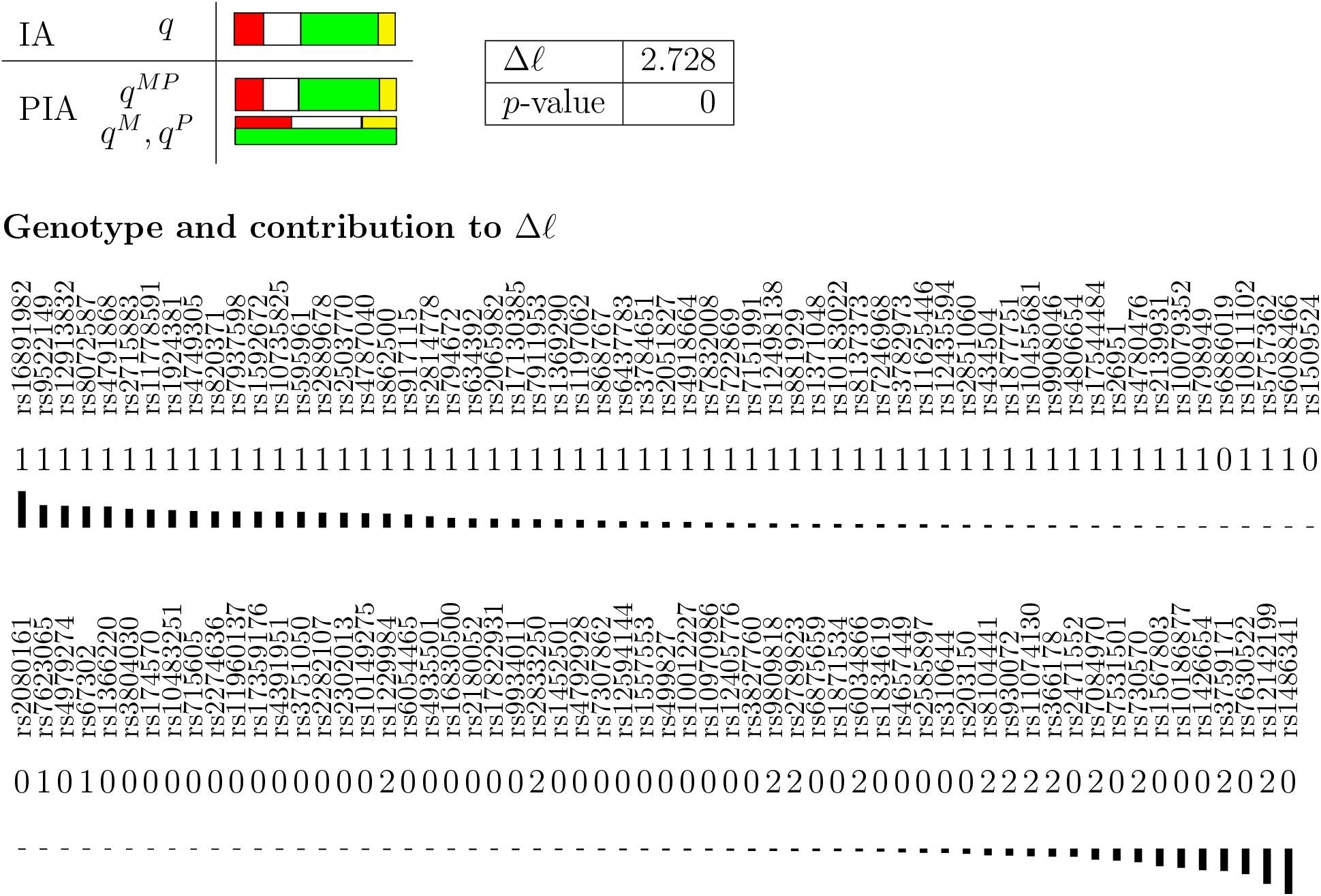

##### Kidd AIMset

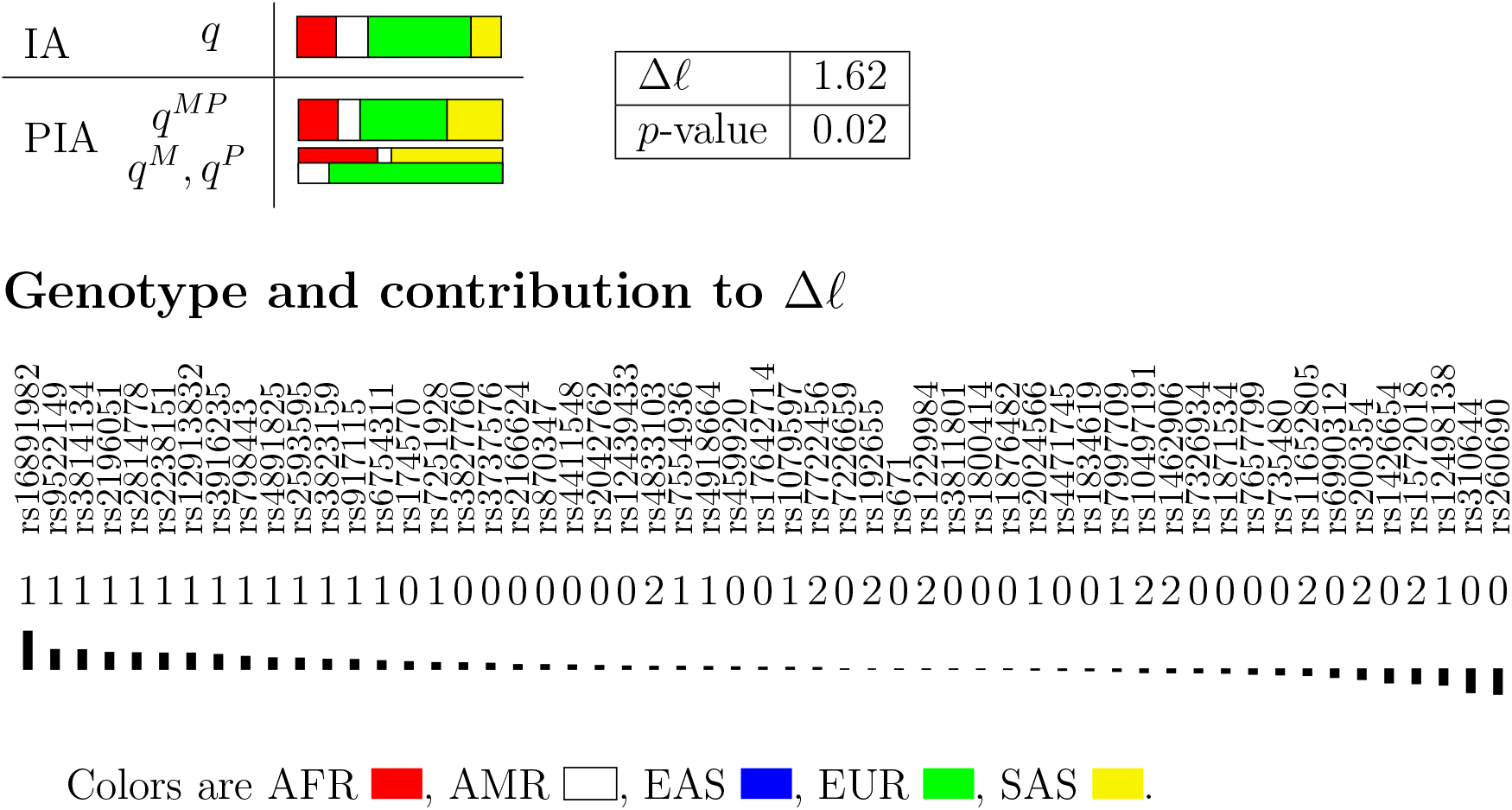

#### Analysis of HG01149, (AMR, CLM)

##### EUROFORGEN AIMset

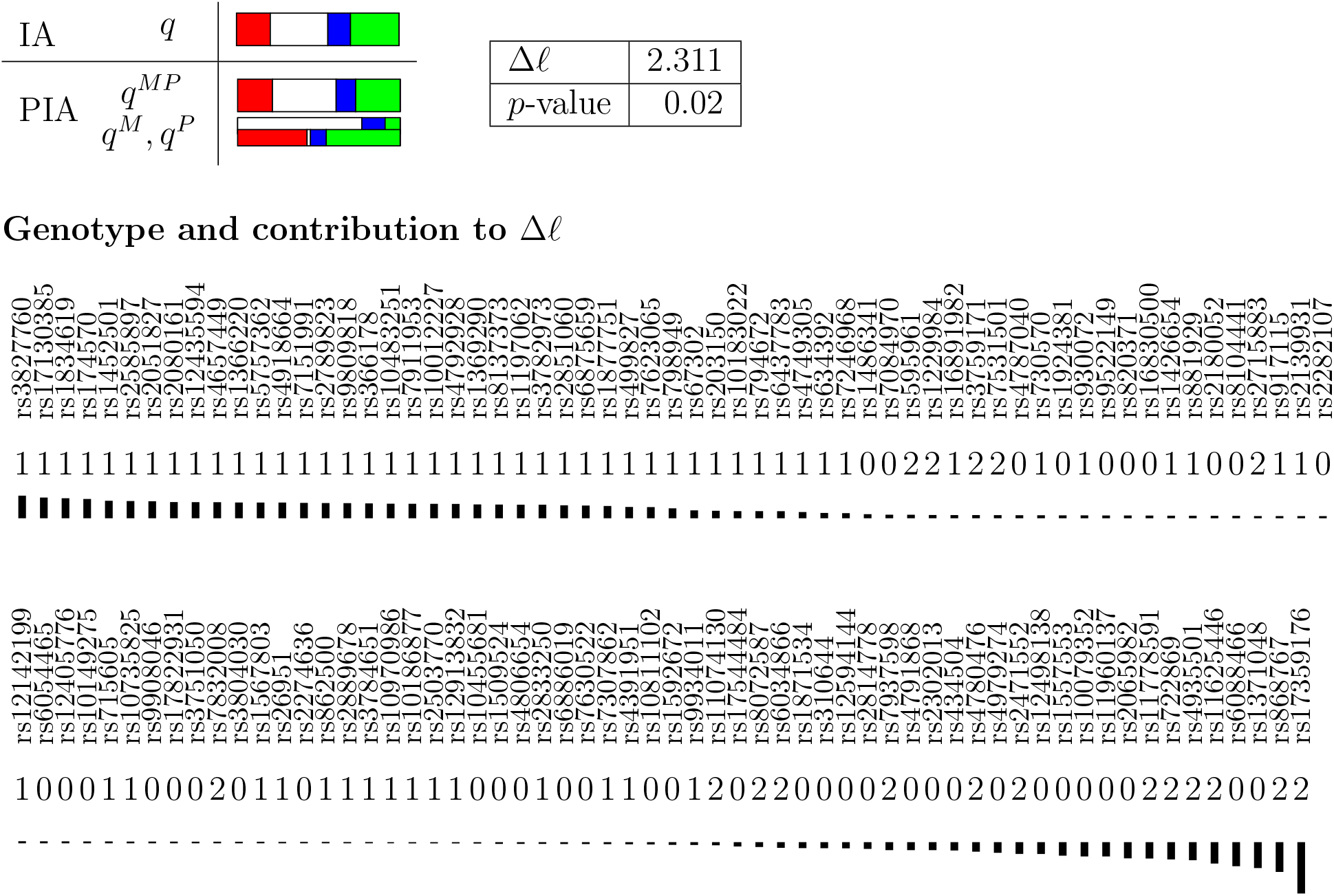

##### Kidd AIMset

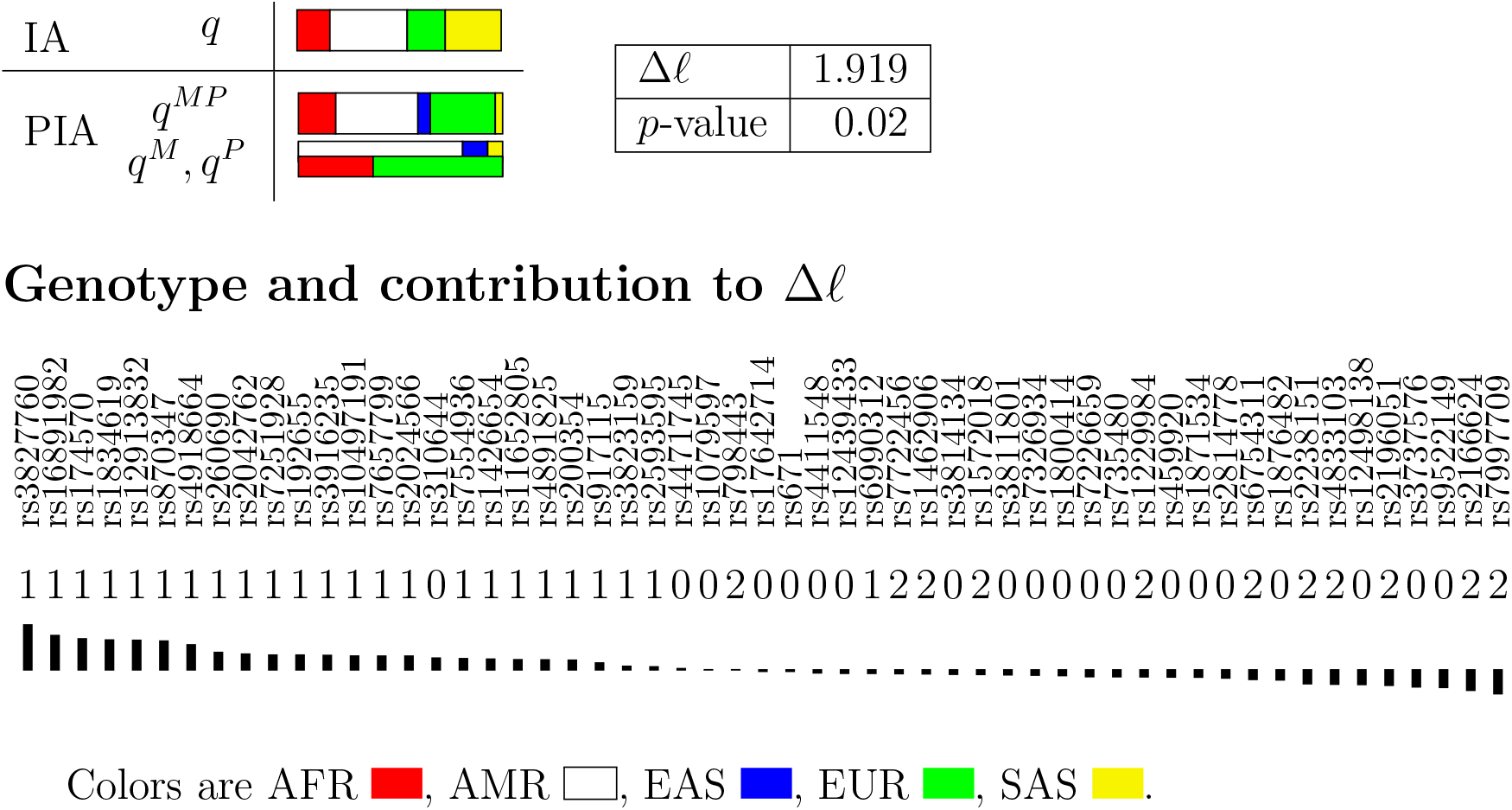

#### Analysis of HG01383, (AMR, CLM)

##### EUROFORGEN AIMset

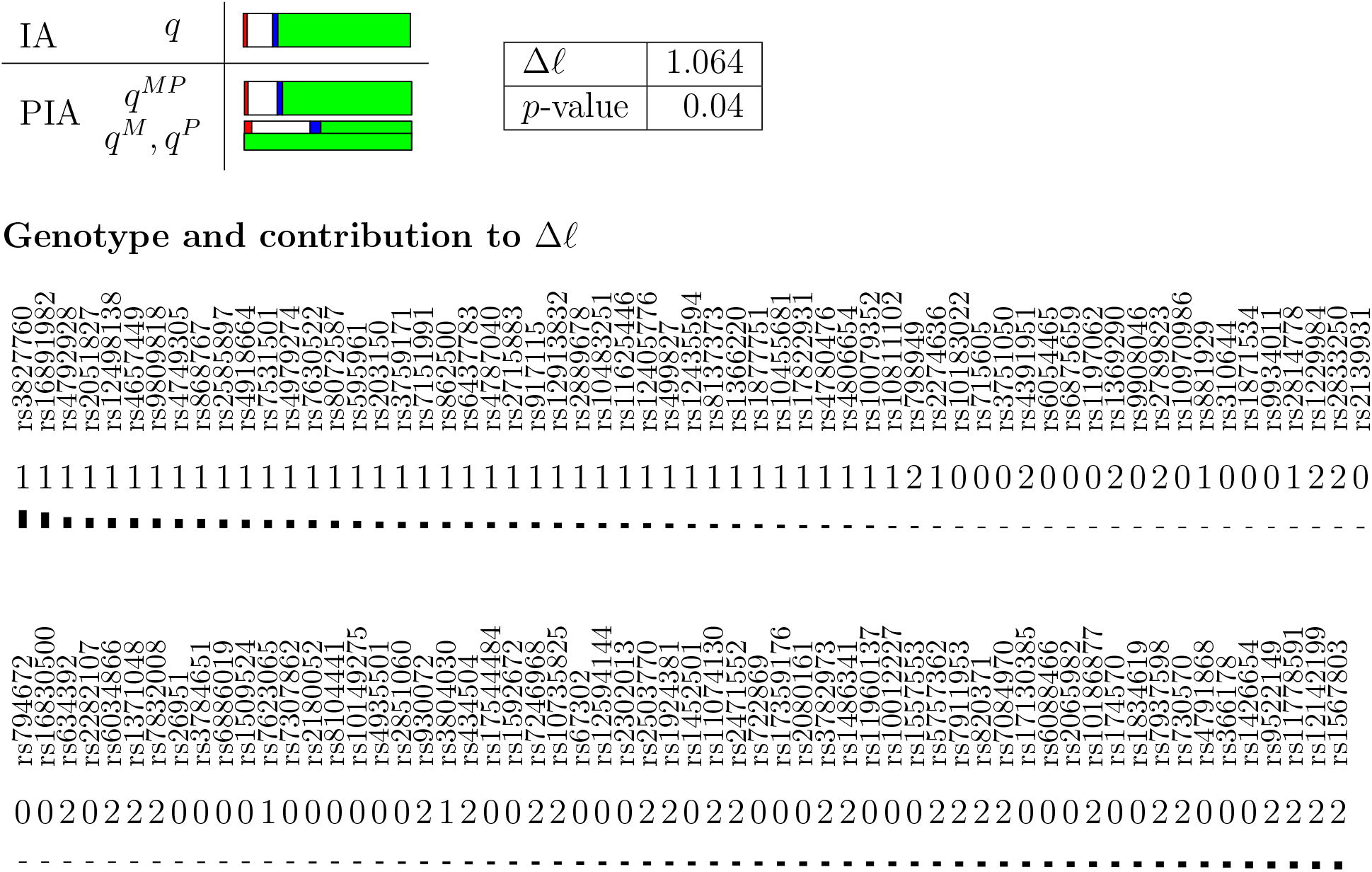

##### Kidd AIMset

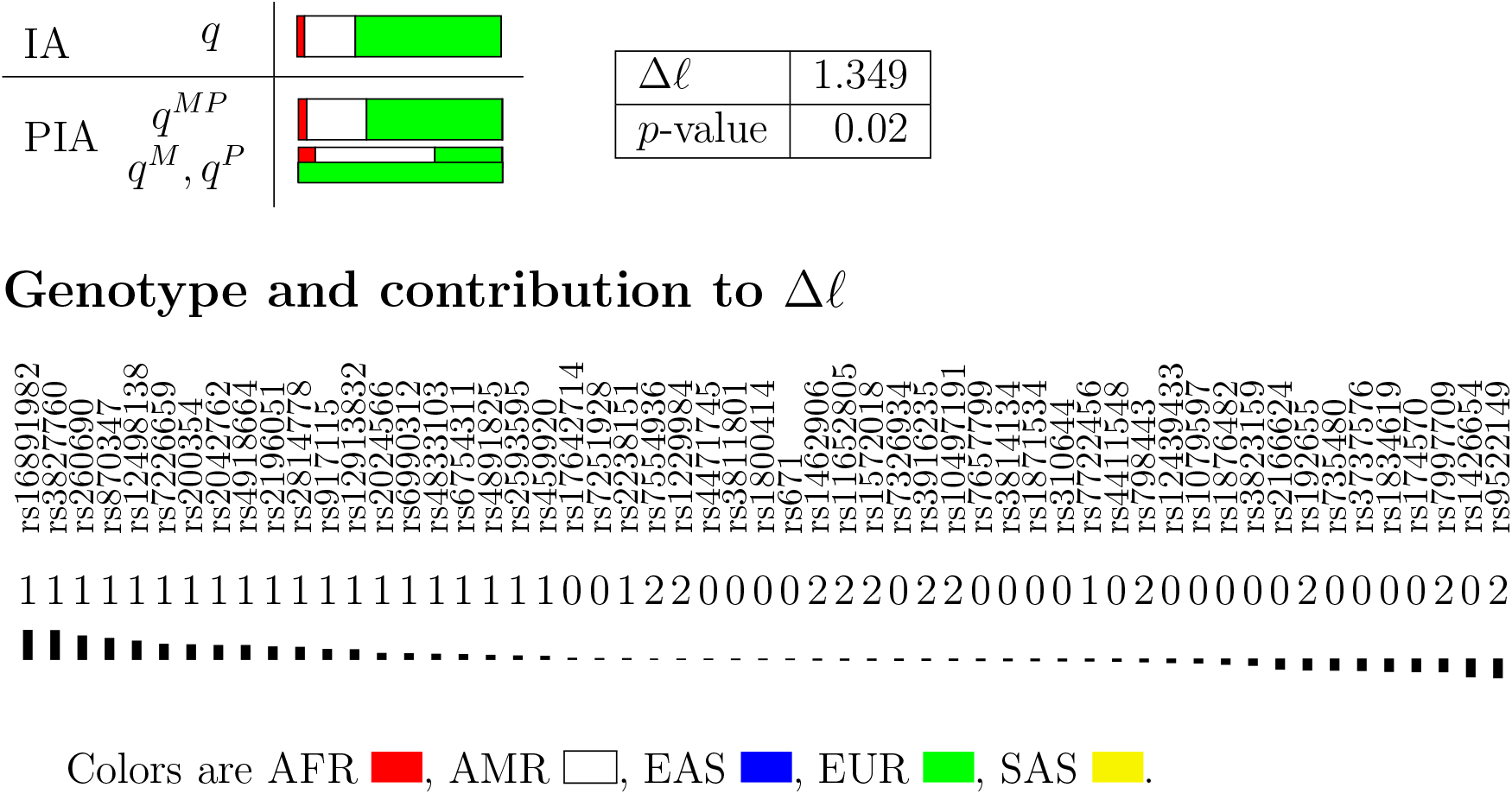

#### Analysis of NA19625, (AFR, ASW)

##### EUROFORGEN AIMset

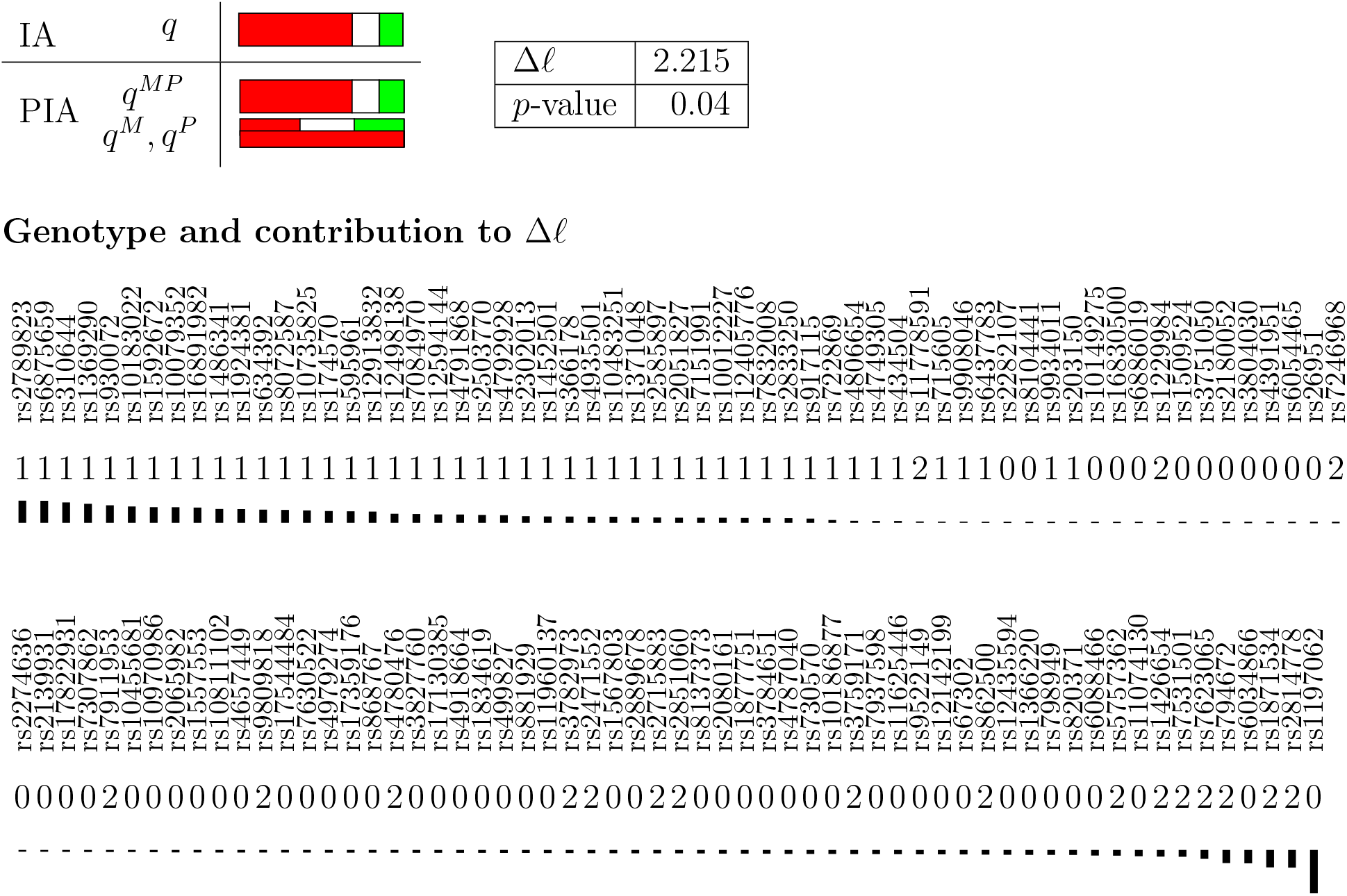

##### Kidd AIMset

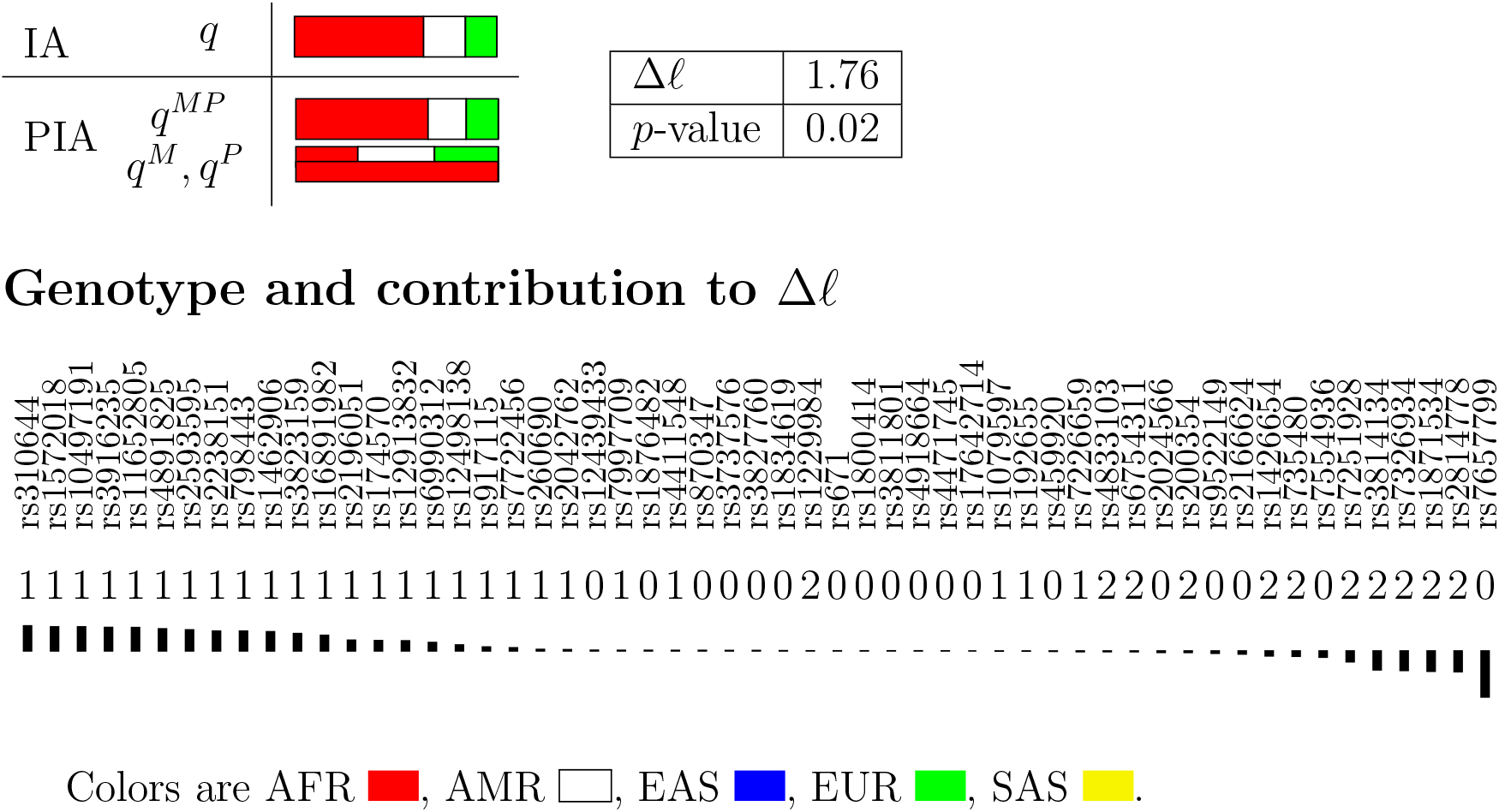

#### Analysis of NA19720, (AMR, MXL)

##### EUROFORGEN AIMset

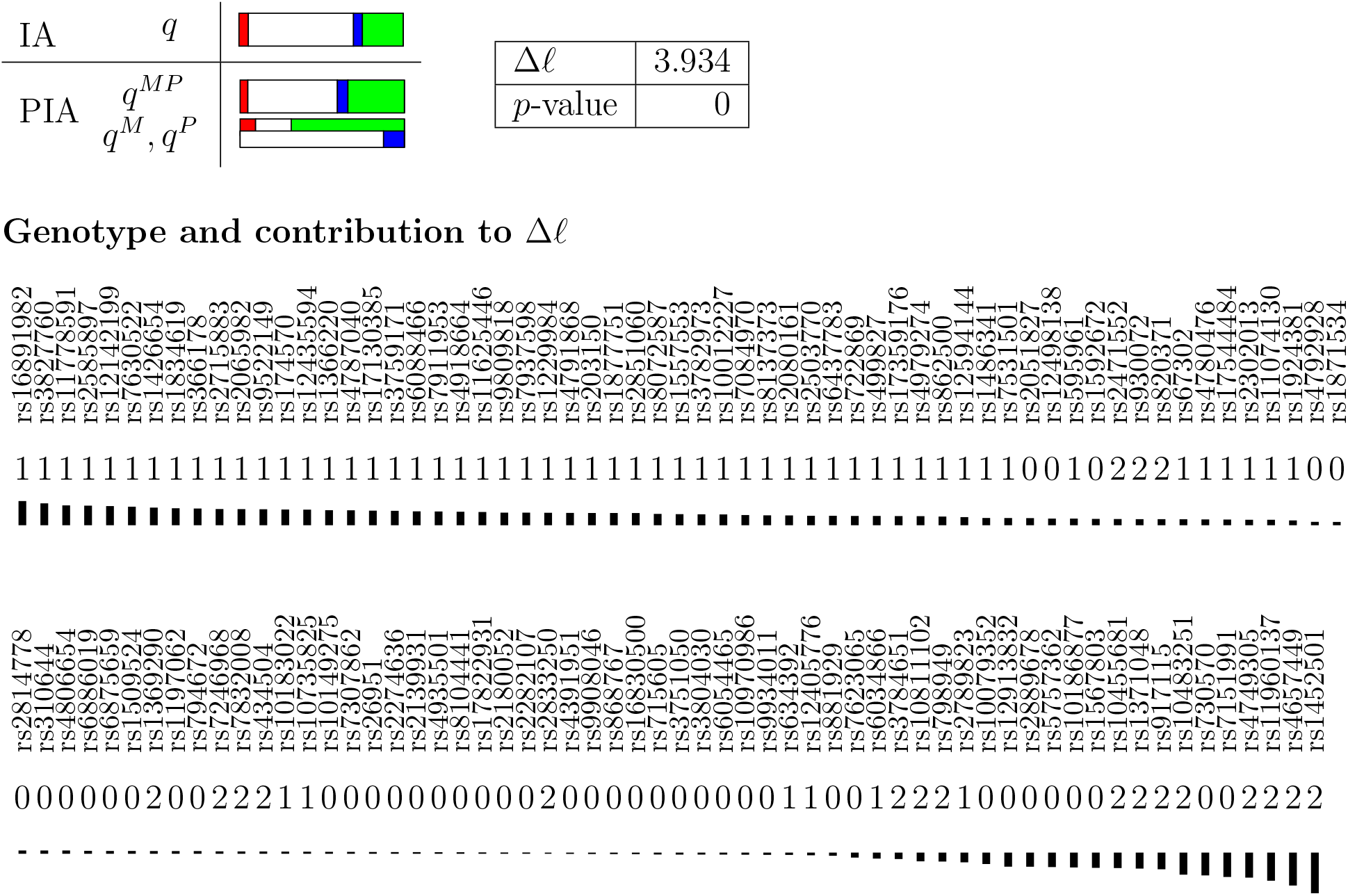

##### Kidd AIMset

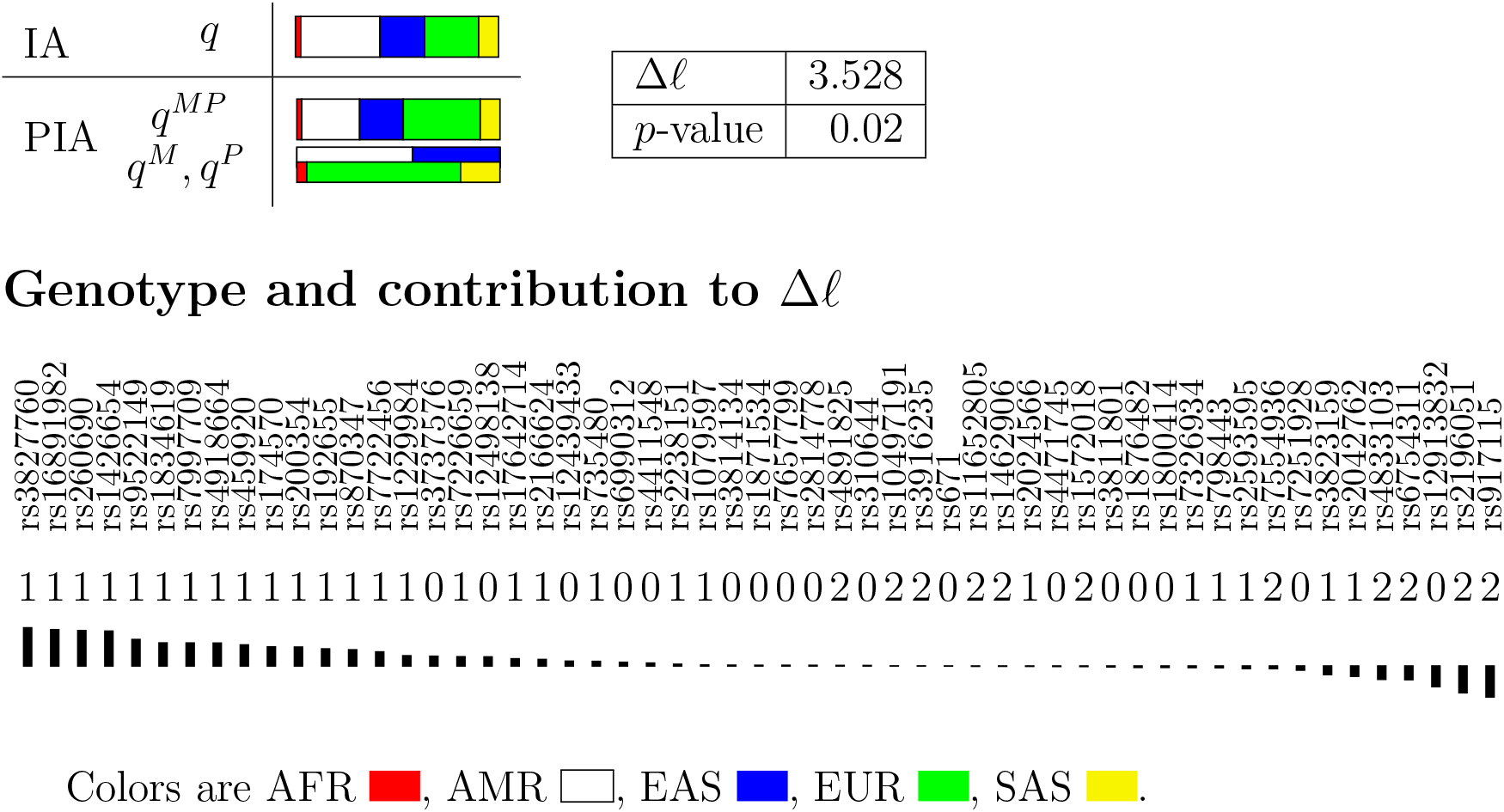

#### Analysis of NA20274, (AFR, ASW)

##### EUROFORGEN AIMset

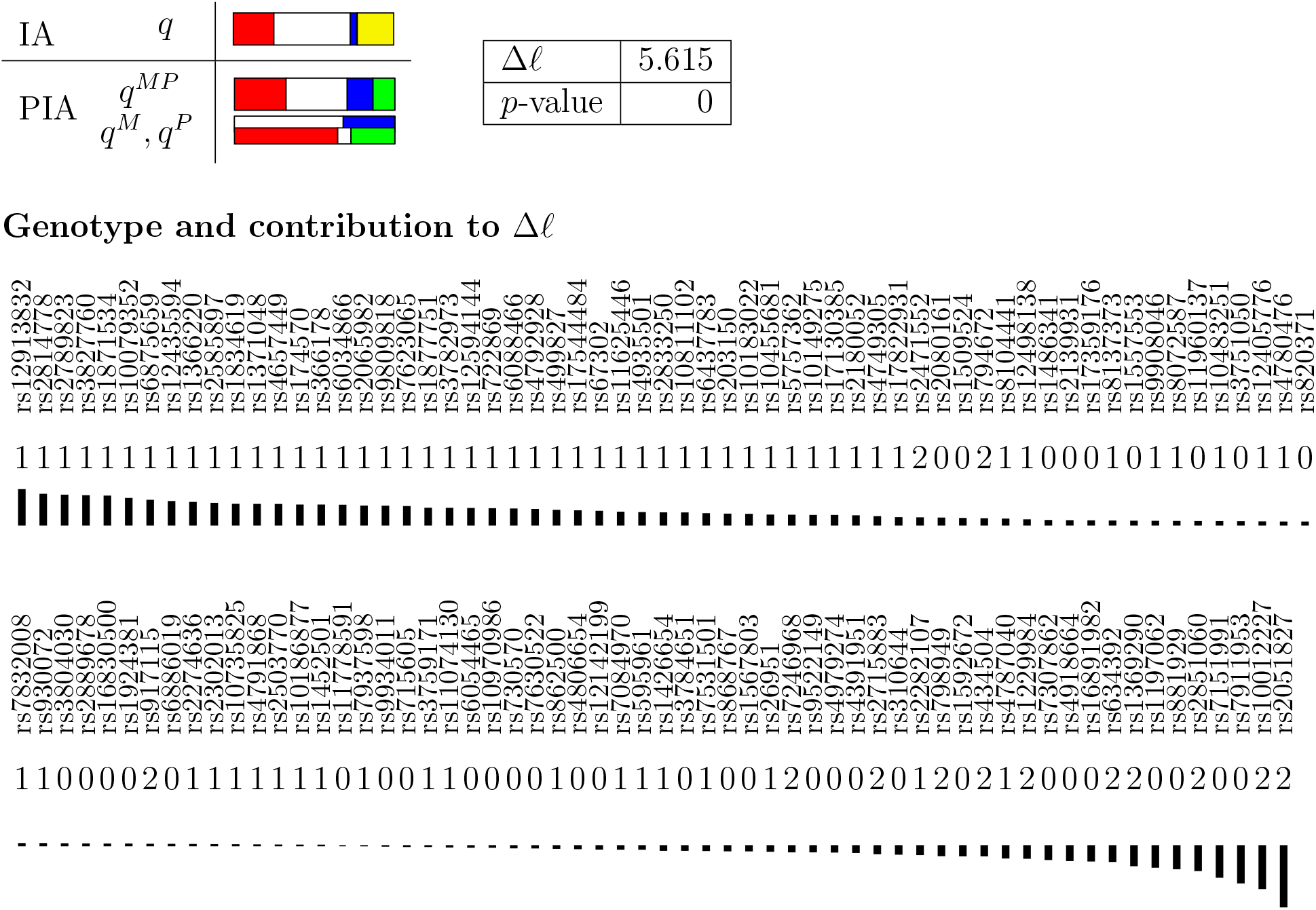

##### Kidd AIMset

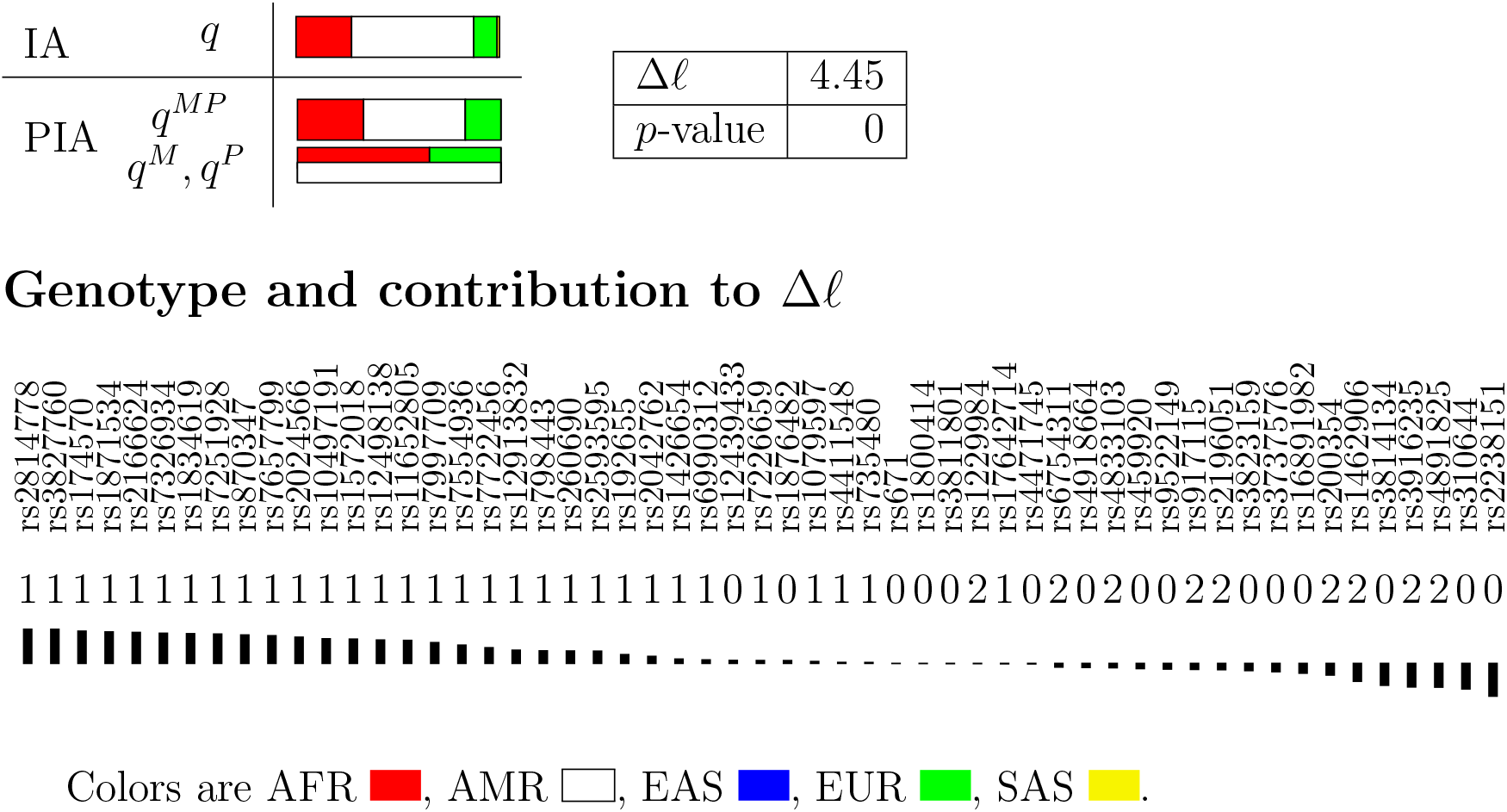

#### Analysis of NA20278, (AFR, ASW)

##### EUROFORGEN AIMset

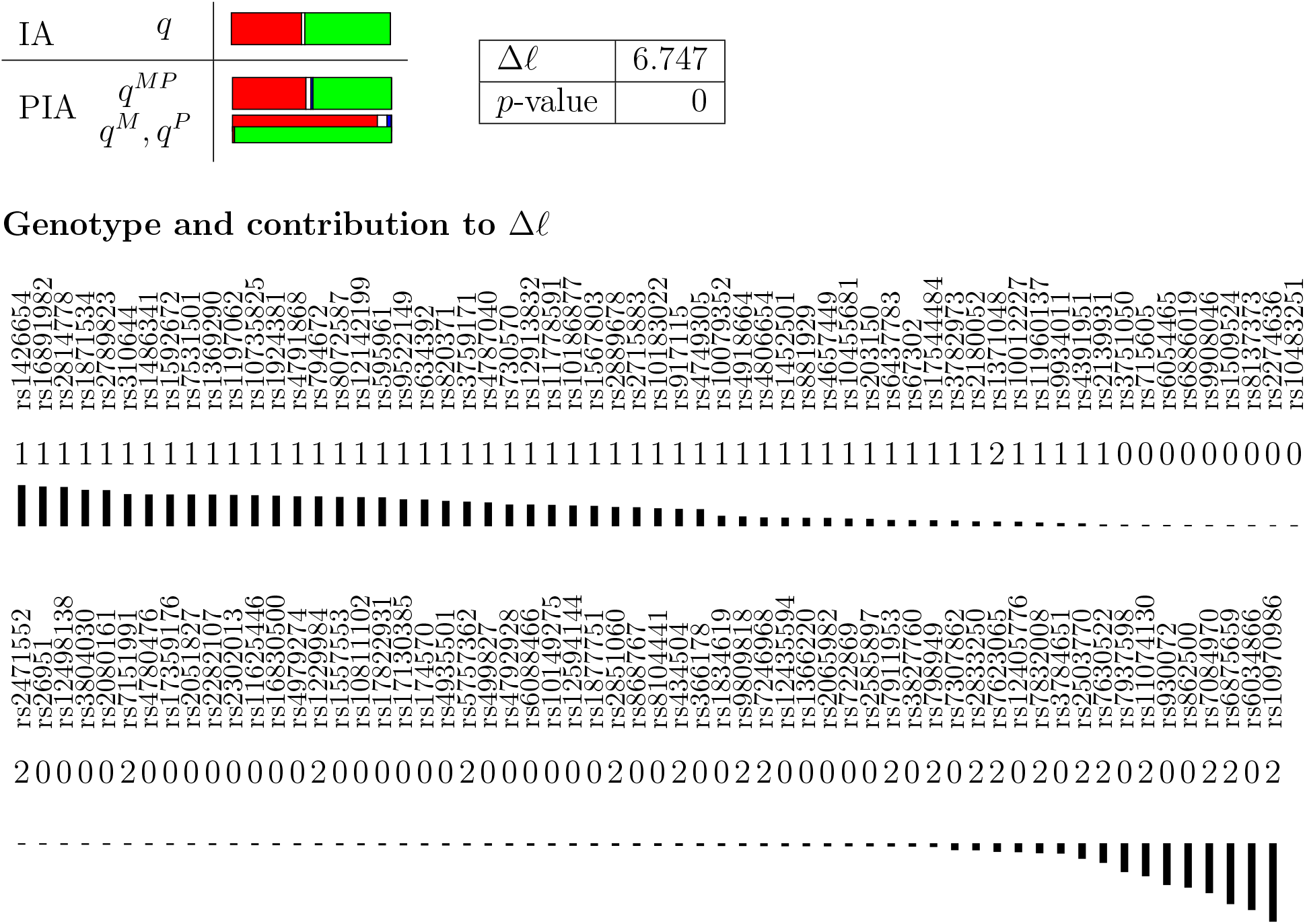

##### Kidd AIMset

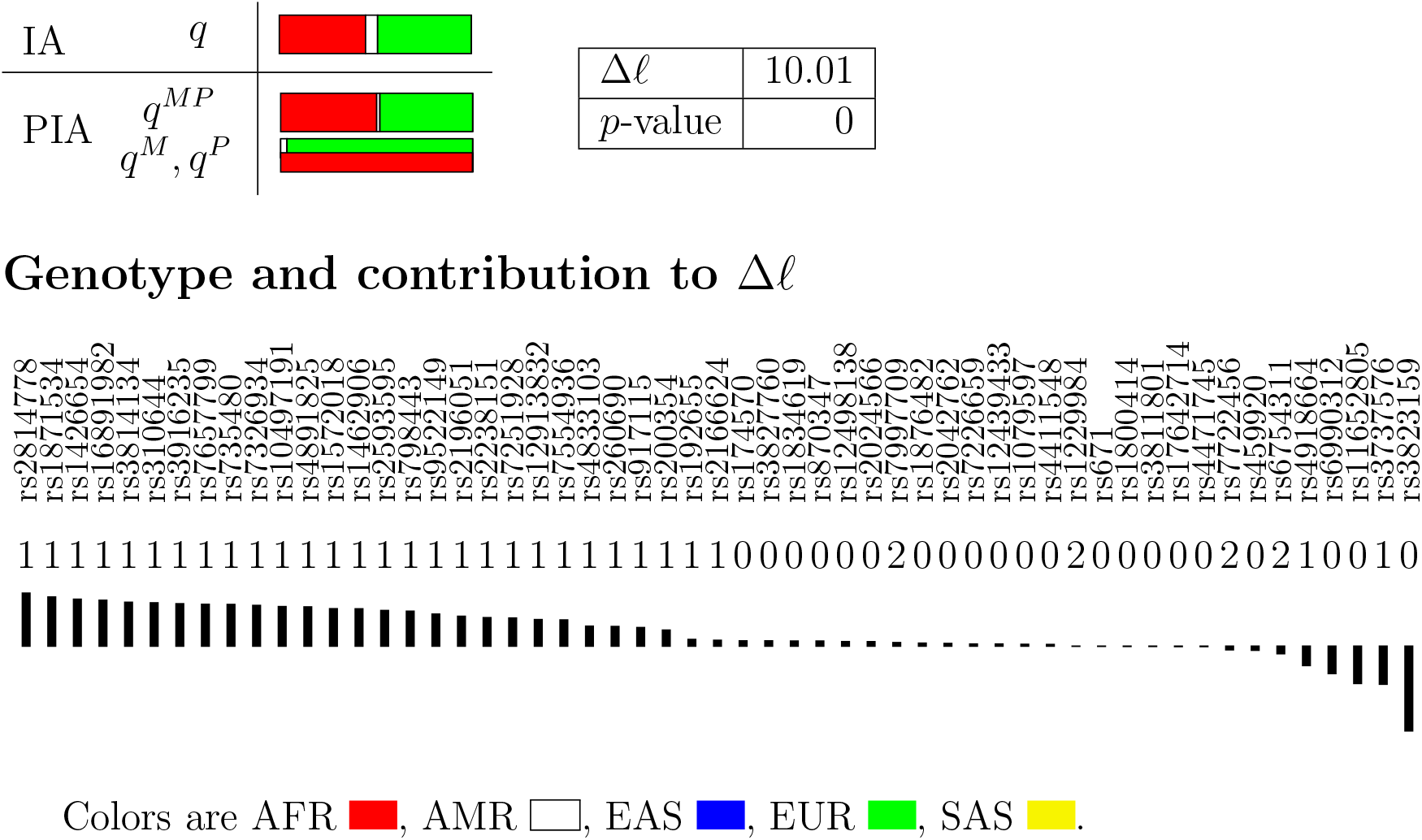

#### Analysis of NA20342, (AFR, ASW)

##### EUROFORGEN AIMset

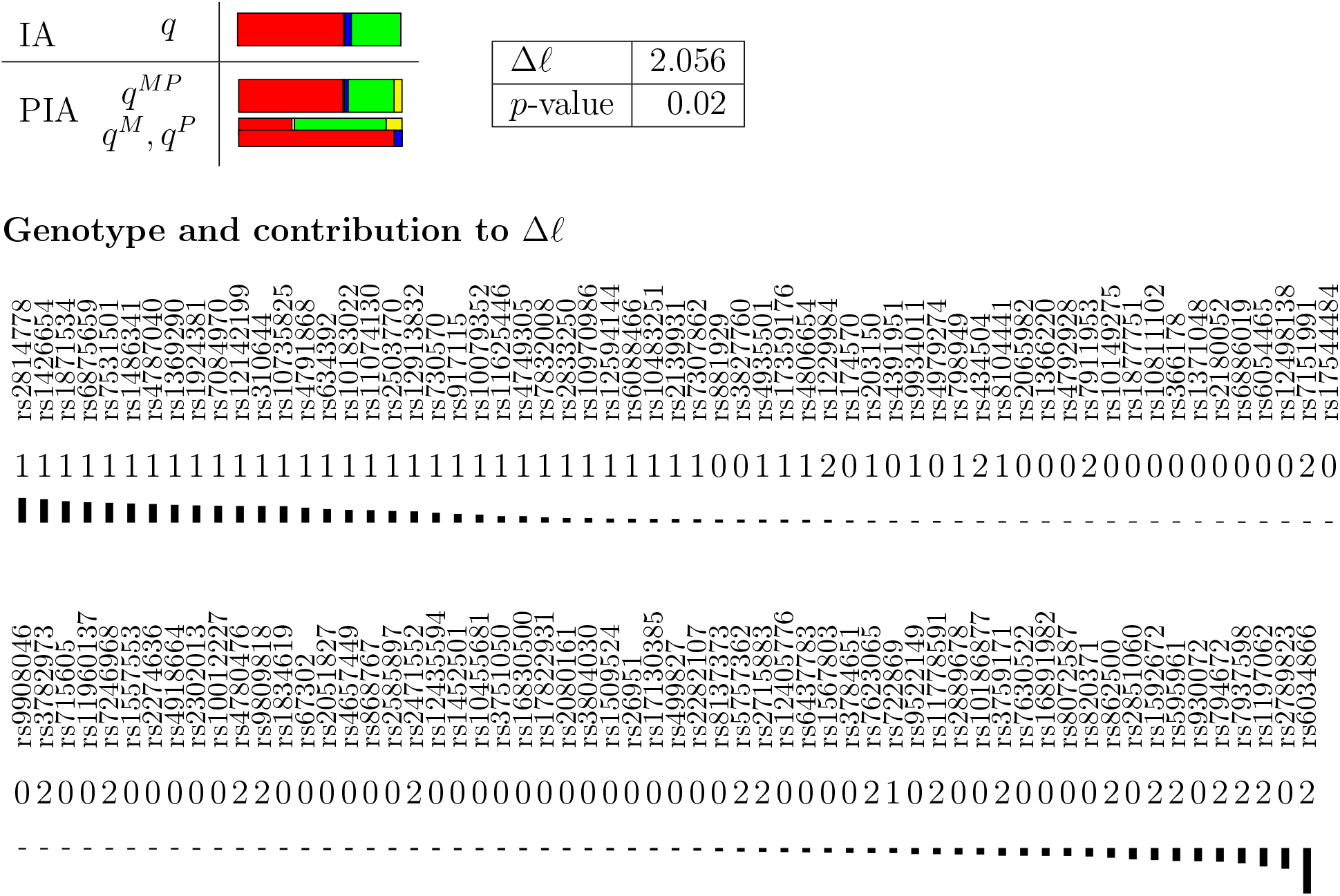

##### Kidd AIMset

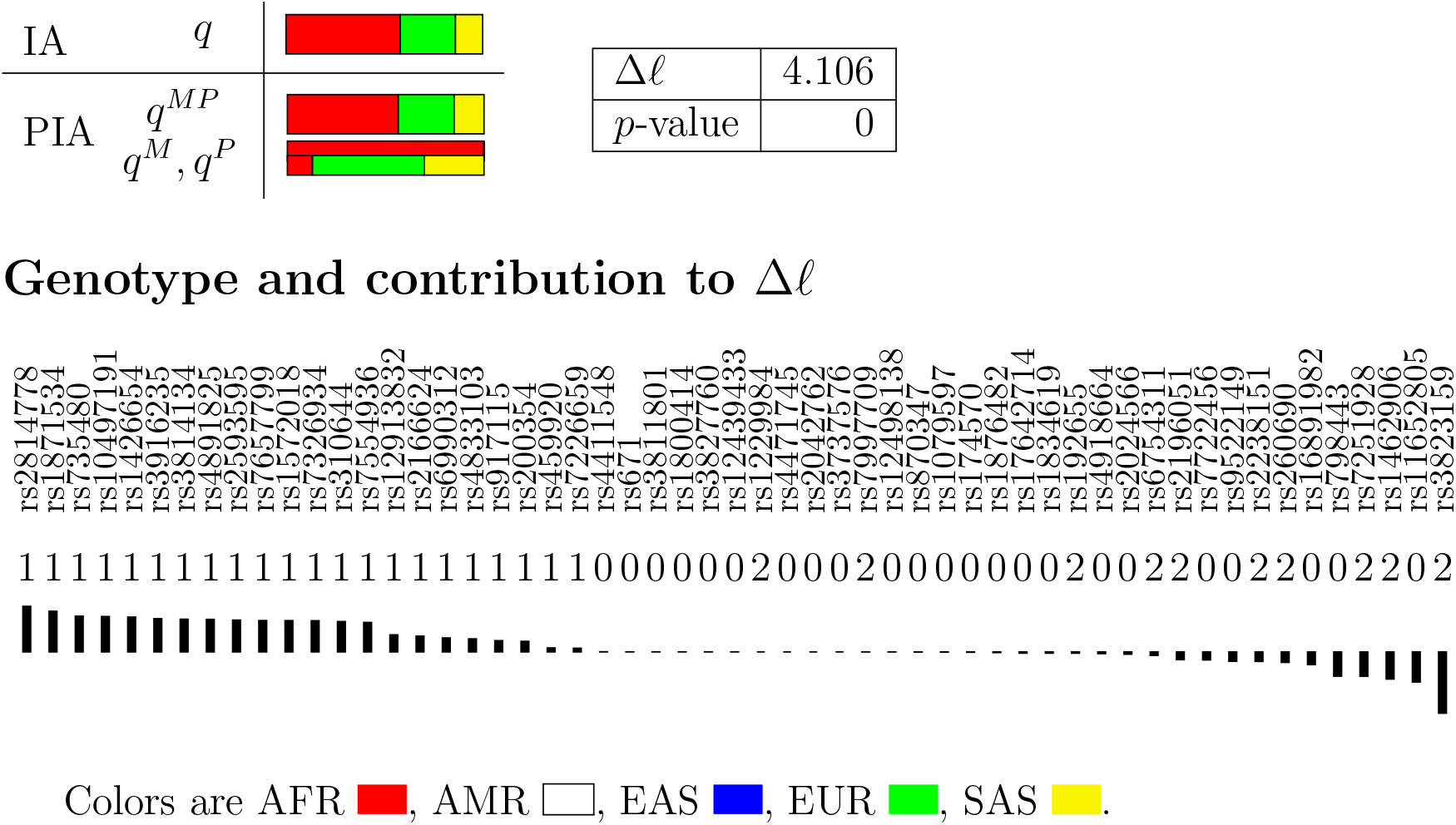

### S2.6 Recent admixture of self-collected samples

#### Analysis of DNA19-1311-83

##### Self description

Mother: Philippines; Father: Germany.

##### Illumina AIMset

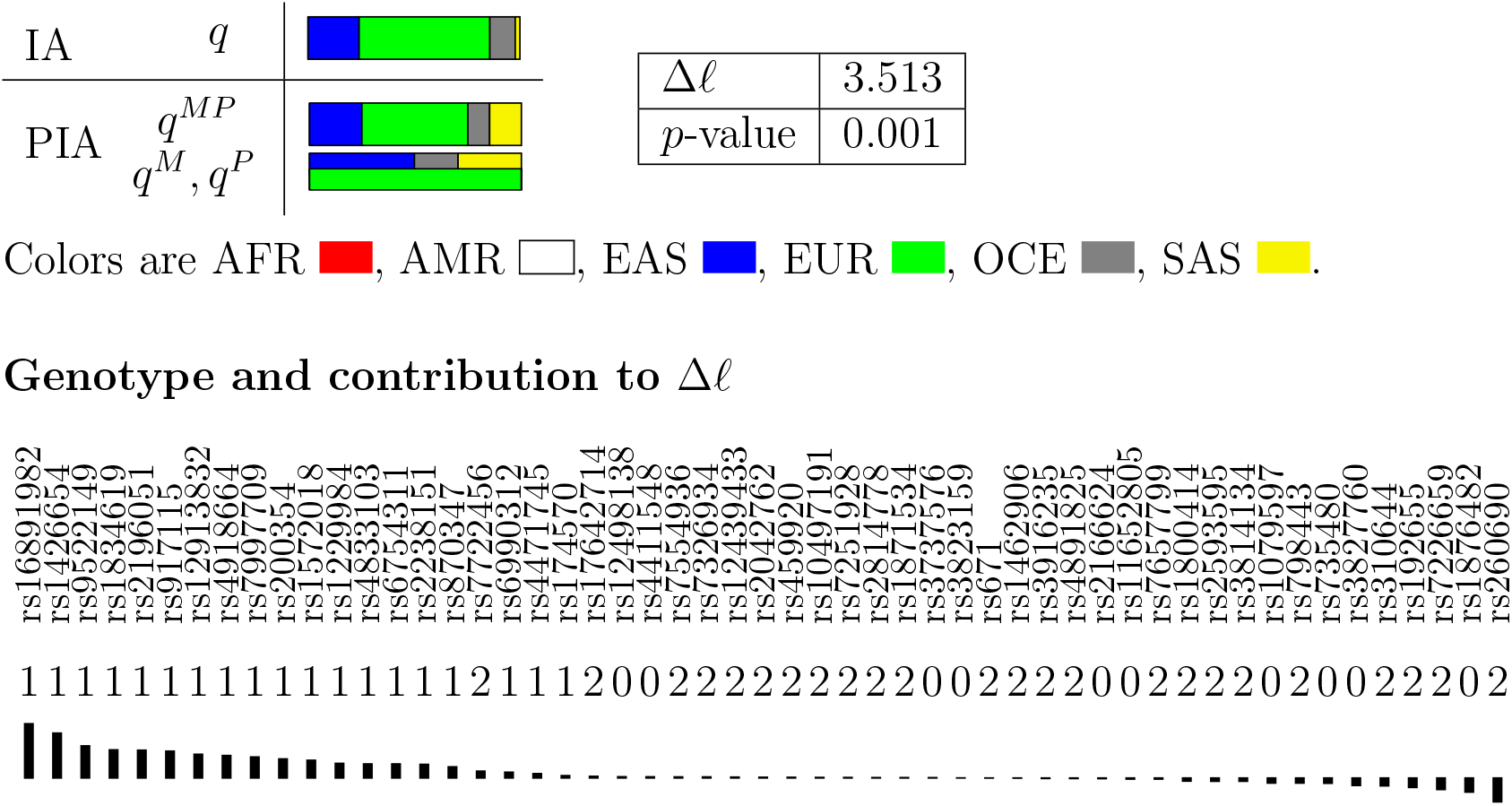

#### Analysis of DNA19-1311-107

##### Self description

Mother: Venezuela; Father: Italy.

##### Illumina AIMset

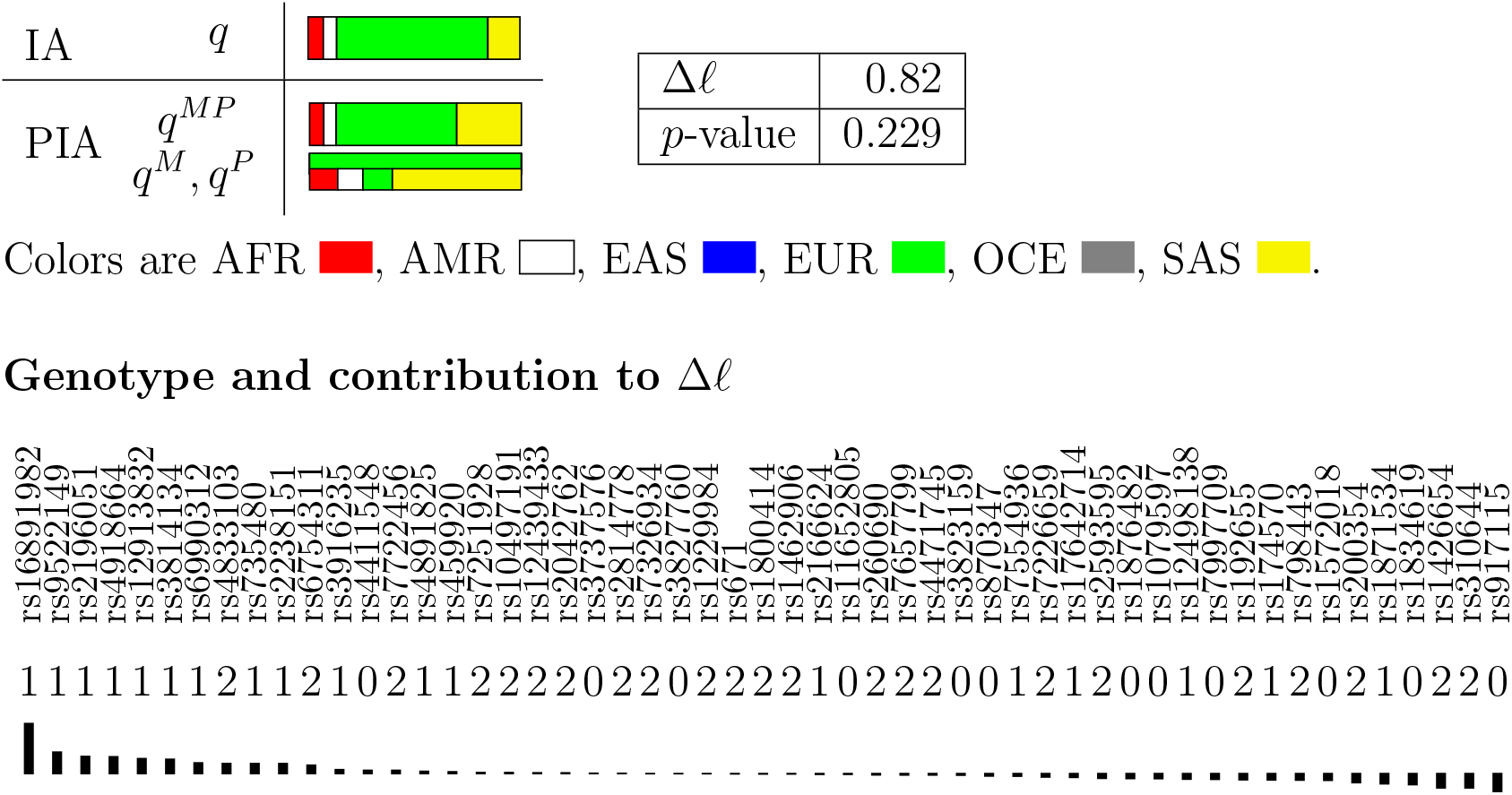

